# Quantitative evaluation of the role of the microenvironment in the phenotype, metabolism, and drug resistance of PDAC tumor organoids

**DOI:** 10.1101/2024.12.31.630904

**Authors:** Ioritz Sorzabal, Xabier Morales, Iván Cortés-Domínguez, Maider Esparza, Lucía Grande, Pedro Castillo, Shruthi Narayanan, Mariano Ponz-Sarvise, Silvestre Vicent, Carlos Ortiz-de-Solórzano

**Author notes:** These authors contributed equally to this work (Ioritz Sorzabal, Xabier Morales, Iván Cortés-Domínguez).

## Abstract

Tumor organoids are powerful tools to study cancer. However, the clinical translation of organoid-based studies depends on how closely the organoid scaffolding material recapitulates the tumor extracellular matrix (ECM). We present a quantitative analysis of the effect of scaffold composition on the phenotype, tissue remodeling, metabolism, and drug resistance of pancreatic ductal adenocarcinoma (PDAC) organoids. We grew PDAC organoids within hydrogels made of Matrigel, collagen-I, or a mixture of collagen-I and Matrigel. Our results show that 1) PDAC organoids grown in Matrigel-only are phenotypically and metabolically similar to low-physiological two-dimensional PDAC cultures; 2) collagen-containing hydrogels, and especially so those grown in hydrogels of mixed collagen-Matrigel composition, accurately reproduce invasive tumors that underwent epithelial-to-mesenchymal transition (EMT) regarding phenotype, tissue remodeling, metabolic activity, and drug resistance. In summary, our work illustrates the importance of three-dimensional (3D) matrix composition for PDAC growth and postulates that organoids grown in collagen-Matrigel hydrogels are the most adequate for studying the biology of invasive PDAC tumors.

## Introduction

Pancreatic cancer is the fourth leading cause of cancer-related deaths, with a five-year survival rate below 11%. PDAC, a pan-exocrine tumor, accounts for 90% of all pancreatic cancer cases (Sally et al., 2022). According to the American Cancer Society, PDAC-related deaths will become one of the top causes of cancer mortality by 2030 (Nakaoka et al., 2023), in part due to its late-stage diagnosis and the limited efficacy of standard chemotherapy options (Chan et al., 2016; Hosein et al., 2022). Histologically, PDAC tumors are highly desmoplastic, as the native ECM is replaced by a dense network of collagen-I, -III, and -IV, laminin, and fibronectin. This rigid tumor microenvironment (TME) promotes the migration and invasiveness of PDAC cells and creates a physical barrier that limits the penetration of therapeutic drugs and immune cells (Ferrara et al., 2021). Thus, modeling this complex tumor-to-TME interaction is of paramount importance.

A deeper understanding of PDAC biology is essential for improving patient prognosis (Hosein et al., 2022). This requires advancing research models that more accurately reflect the complex PDAC tumor physiopathology. Traditional two-dimensional cell models cannot replicate the full range of environmental cues found *in vivo*, especially so regarding the presence of realistic cell-cell and cell-ECM interactions (Bahcecioglu et al., 2020). Animal models provide a closer approximation to PDAC’s physiopathology. However, they are costly, allow limited experimental flexibility, and raise significant ethical concerns related to the number of animals needed and the level of suffering inflicted on them (Abdolahi et al., 2022; Chang & Lin, 2021). These limitations underscore the need for novel, physiologically relevant models with high translational potential.

Organoid models, derived from tumor samples grown in 3D ECM biomimetic lattices, have become a promising alternative to traditional models (LeSavage et al., 2022). Tumor organoids retain key genetic, proteomic, and morphological features of the original tumors, including their unique intra- and inter-tumoral heterogeneity. Patient-derived tumor organoids (PDOs), established from clinical samples, have direct translational value (Thorel et al., 2024; Yang & Yu, 2023), but their use faces challenges related to availability and reproducibility. However, organoids derived from murine cancer models representative of the human disease (Boj et al., 2015) provide greater availability and experimental flexibility (Huang et al., 2021; Marshall & Mason, 2019) and may fill the translational gap derived from PDOs scarcity, thus becoming ideal platforms for exploring cancer initiation, tumor-ECM interactions, and cancer metastasis (W. Li et al., 2023). However, the design of the biomimetic scaffolds that support these organoids, sometimes simplified, unreported, or overlooked, is crucial, as it must accurately replicate the mechanobiology of TME (Driehuis et al., 2020).

The TME is a highly complex ecosystem consisting of vasculature, immune cells, and the acellular ECM (Giraldo et al., 2019), which comprises a highly organized 3D network of fibrous proteins (e.g., collagen, fibronectin, laminin), soluble signaling factors, bioactive molecules, and mechanotransduction cues (Morales et al., 2021; Sullivan et al., 2022). In healthy tissues, the TME is tightly regulated to maintain homeostasis. During cancer progression, this delicate balance becomes disrupted, leading to abnormal interactions between tumor cells and the surrounding ECM that trigger changes in the tumor phenotype, which in turn alter the biomechanical properties of the ECM (Morales et al., 2021; Sainio & Järveläinen, 2020). These changes contribute to tumor progression, often through key pathways like the transforming growth factor (TGF)-β signaling pathway (Sainio & Järveläinen, 2020; Yuan et al., 2023). In PDAC tumors, the ECM undergoes extensive remodeling caused by the deposition of newly formed fibers and the crosslinking and alignment of the existing ones. These structural changes enhance tumor cell migration and invasiveness (Anguiano et al., 2020) and promote malignant cell transformation through the emergence of mesenchymal-like traits (Rice et al., 2017), which have been shown to be promoted by TGFβ in PDAC (Principe et al., 2021). These insights emphasize the crucial role of the ECM in tumor development and drug resistance. Therefore, the ECM’s mechanobiological properties must be carefully considered when developing realistic organoid models. Indeed, the biomimetic scaffolds used to sustain the organoids must accurately replicate the ECM properties. In contrast, ‘simplified’ hydrogels produce organoids with phenotypic characteristics that diverge from native tumors (Heo et al., 2022; Sullivan et al., 2022).

Matrigel, derived from the TME of the Engelbreth-Holm-Swarm mouse sarcoma, is the most used scaffold for organoid culture. Matrigel is a rich source of ECM components, including collagen-IV, laminin and, enactin, along with a variety of bioactive cues. However, its use has some drawbacks, including batch-to-batch variability, mechanical softness, and lack of fibrous structure, which negatively affect experimental reproducibility and limit the physiological relevance of the models. Synthetic hydrogels (Kozlowski et al., 2021) based on materials such as Polyethylene-Glycol (PEG), Polycaprolactone (PCL), Poly(lactic-co-glycolic) acid (PLGA), or Polyisocyanide (PIC) derivatives, offer optimal tunable structural and biomechanical properties and are more reproducible than Matrigel-based hydrogels (Gan et al., 2023) being ideal to study the effect of isolated biomechanical cues but not to mimic the complexity of the native ECM, as these materials typically lack cell binding sites and require bio-functionalization to enhance cell adhesion. Furthermore, they require the addition of complex growth factor mixtures to approximate the rich ECM of the tumors. Finally, hydrogels derived from natural polymers, such as collagen-I and fibrin proteins, or polysaccharides like hyaluronic acid and alginate, have been widely used to culture organoid models. These polymers are inherently bio-functional but have suboptimal rheological properties, as a complicated equilibrium in the concentration of the protein is crucial for allowing efficient handling of the pre-polymerized protein solution while ensuring a polymerized hydrogel with sufficient structural stability (Heo et al., 2022).

In this study, we present a comprehensive analysis of the use of biomimetic hydrogels made of Matrigel, collagen-I, or a collagen-Matrigel mixture (Anguiano et al., 2017, 2020) for PDAC organoid development. Collagen-I ensures biofunctionality and modulates the matrix stiffness, helping recapitulate the fibrotic ECM of PDAC tumors (Ferrara et al., 2021). Meanwhile, Matrigel provides a rich biochemical environment made of proteins commonly found in the ECM and modulates the viscous component of the material (Chaudhuri et al., 2020; L. Li et al., 2024). We evaluated the effect of the composition of these hydrogels on several phenotypic descriptors, including “early-stage organoid seeds” (EOSs) migration, mature organoid morphology, matrix remodeling, metabolic reprogramming, and the expression of epithelial to mesenchymal transition (EMT) markers under regular media or TGFβ conditioning. Furthermore, we evaluated the impact of the hydrogel biomechanical properties on the resistance of PDAC organoids to Gemcitabine (Gem), both *in vitro* and *in vivo* subcutaneously implanted in an immunocompetent syngeneic murine model. Our results reveal significant differences in the phenotype of PDAC organoids as a function of hydrogel composition, underscoring the essential role of ECM mechanobiology in the development and chemoresistance of PDAC organoids, and offer valuable guidance for researchers using these models, especially in drug discovery assays and in the clinical translation of pancreatic cancer biology studies.

## Results

### Mechanical characterization of the biomimetic hydrogels

To contribute to understanding the role of the ECM in PDAC progression, we explored the effect of the matrix composition in the biology of PDAC organoids. We used three hydrogels of distinct morphological and biomechanical properties: Matrigel only (M, 4 mg/mL); collagen-only (C, 4 mg/mL); and a mixture of collagen-Matrigel (CM, 2 mg/mL-2 mg/ml). We first characterized the rheology and microstructure of the hydrogels: The rigidity and elastic component of the scaffolds, associated with the storage modulus (G’), increases with the increasing concentration of collagen-I and decreasing concentration of Matrigel (**Supplementary Figure 3A, C**). A similar trend was found for the viscosity, associated with the loss modulus (G’’) (**Supplementary Figure 3B, D**) The ratio between G’’ and G’ (TanΔ) tends to zero in all three hydrogels, revealing a predominantly solid, elastic behavior (**Supplementary Figure 3E**), and increases as Matrigel is replaced by collagen-I, indicating a higher influence of the viscous fraction of the material due to the presence of Matrigel. Second Harmonic Generation (SHG) imaging of collagen-I fiber distribution in C and CM hydrogels (**Supplementary Figure 3F**) showed no significant differences in collagen-I fiber thickness caused by the presence of Matrigel (**Supplementary Figure 3G**). However, mixed collagen-Matrigel matrices displayed larger pore size diameter (**Supplementary Figure 3H**) and lower fiber density (**Supplementary Figure 3I**) compared to C hydrogels.

### Collagen-I and TFGβ promote the invasiveness of early-stage PDAC organoid seeds

Prior to organoid formation, PDAC cells actively migrate through the hydrogel, establishing cell-to-cell contacts and clustering into EOSs. We analyzed the migration of EOSs in time-lapse microscopy videos (**Supplementary Video 1**). Representative static images and track plots are shown in **Figure 1A, B**. The results of the quantification show a significant increase in the mean migration speed (**Figure 1C**), as well as in the number and length of cell protrusions of the EOSs (**Figure 1E, F**), between hydrogels M and CM, and between CM and C hydrogels. Furthermore, TGFβ conditioning enhanced migration speed and the number and length of cell protrusions in all three hydrogels, with greater intensity in those scaffolds containing collagen-I (C, CM). Finally, no significant differences in EOS directness were observed in the 3D cell trajectory plots, displaying a random migration pattern regardless of the matrix composition (**Figure 1D**).

**Figure 1.**
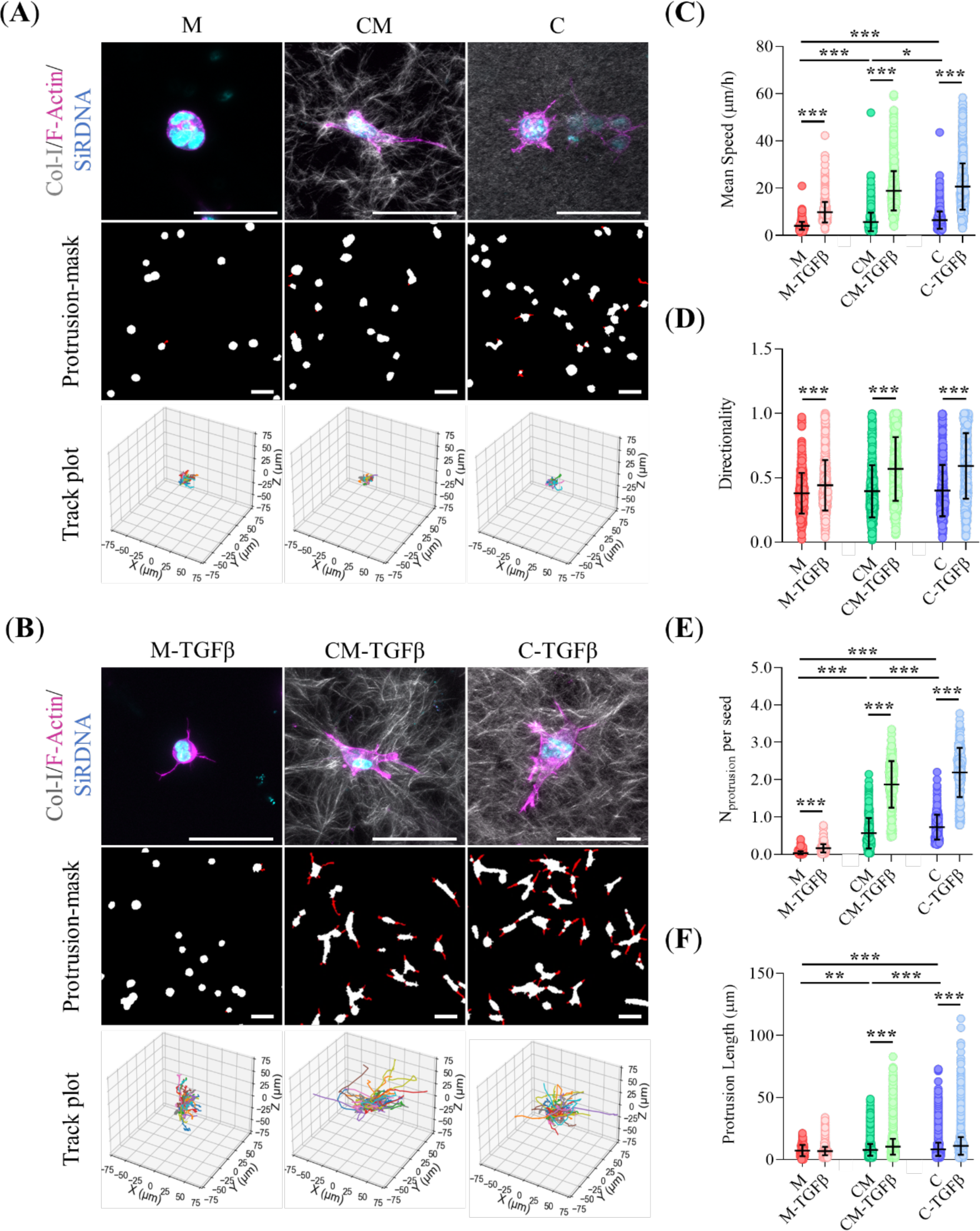
Migration of PDAC EOSs. (**A**-**B**), Z-stack projections of representative 3D CLSM images of EOSs in M, CM, and C hydrogels under regular media or TGFβ conditioning. Staining: F-Actin (magenta), nuclei (SiRDNA, cyan), and the collagen-I lattice (SHG). Protrusion masks (in red) highlight cytoplasm extensions. The 3D track plots illustrate the length and directness of the EOS trajectories. Scale bars: 15 µm (CLSM images), 50 µm (protrusion masks). (**C**-**F**), Quantitative analysis of invasive descriptors extracted from panels A and B included mean speed (μm/h), directionality, and the number and length of protrusions (μm), respectively. *** Indicates highly statistically significant differences (p<0.001). ** Indicates moderately statistically significant differences (p<0.01). * Indicates statistically significant differences (p<0.05).

### The biomechanical properties and TGFβ modulate the morphological diversity of mature PDAC organoids

We then investigated how matrix composition influences the morphology of mature PDAC organoids. We identified three organoid morphologies: cystic, characterized by a prominent lumen surrounded by a ring of cells; solid, consisting of a compact cluster of cells without lumen; and invasive, which includes a variety of compact morphologies with elongated extensions that penetrate the surrounding hydrogel (**Figure 2A-B**). Based on our machine-learning analysis (**Figure 2C**), most organoids grown in Matrigel display cystic morphology (58%), with the remainder being solid (34%) and rare occurrence of invasive forms (8%). Conversely, collagen-containing hydrogels favor the presence of invasive morphologies (53% CM, 46% C) alongside some solid (29% CM, 33% C) and fewer cystic (18% CM, 21% C) morphologies. The presence of TGFβ strongly affected these morphologies, causing an increase in solid forms in Matrigel (91%) and a shift toward invasive morphologies in collagen-containing hydrogels (63% CM-TFGβ, 54% C-TFGβ), the remaining organoids predominantly displaying solid morphologies. Interestingly, TGFβ conditioning significantly decreased the number of organoids across all hydrogels compared to non-TGFβ-treated conditions (p<0.0001) (Data not shown).

**Figure 2.**
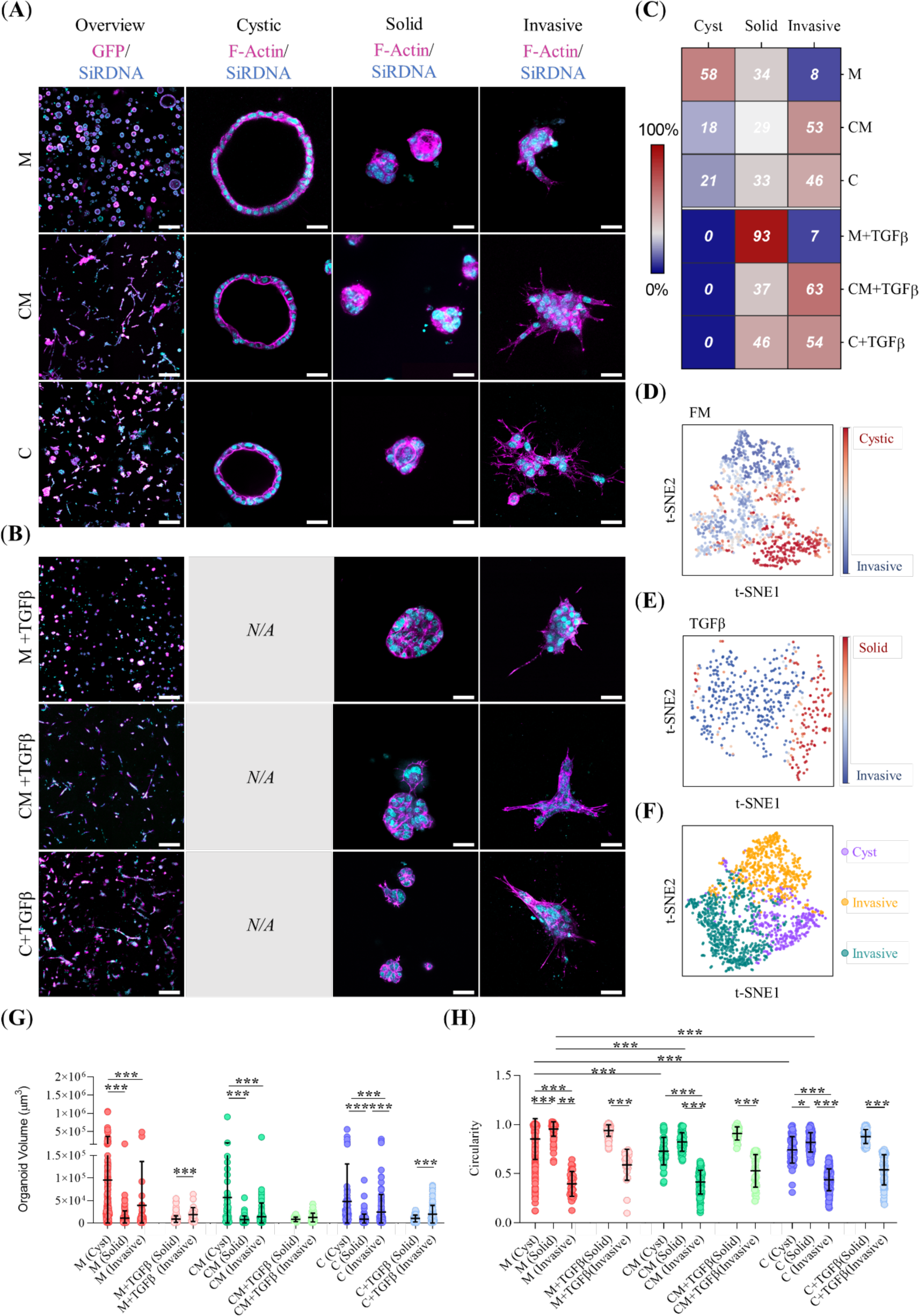
PDAC organoid morphologies. (**A**-**B**), Z-stacks projections of representative 3D CLSM images of GFP-PDAC organoids in M, CM, and C hydrogels under regular media or TGFβ conditioning. Staining: cytoplasm (endogenous-GFP, magenta) and nuclei (SiRDNA, cyan). Low-magnification images (left panel) show the overall organoid morphologies, while high-resolution images (right panel) provide a detailed view of the different organoid morphologies. Scale bars: 150 µm and 25 µm, respectively. (**C**), Abundance heat-map of PDAC organoid phenotypes. (**D**-**E**), t-SNE projection showing the morphological class distribution of different PDAC organoids under regular media or TGFβ conditioning, respectively, combining data from M, CM, and C hydrogels. The scale bar represents the classification probability as Cyst, Solid, or Invasive. (**F**), t-SNE projection showing the distribution of the three morphological classes identified by the SVM classifier, combining data from all hydrogels and media conditions. (**G**-**H**), Quantification of the morphological features used in the SVM classifier, including organoid volume and circularity. *** Indicates highly statistically significant differences (p<0.001). ** Indicates moderately statistically significant differences (p<0.01). * Indicates statistically significant differences (p<0.05).

The morphological diversity of the organoids is further illustrated by the t-SNE plots, which group organoids based on their morphological features. In non-TGFβ-treated hydrogels (**Figure 2D**), cystic and invasive organoids cluster at the upper and lower ends of the t-SNE plot, while solid organoids are positioned in between as a transitional morphology. In TGFβ-treated organoids (**Figure 2E**), solid and invasive forms cluster in two well-defined areas of the t-SNE plot. A more nuanced understanding of the intrinsic abundance of each organoid type was achieved by analyzing the data according to hydrogel type and medium conditioning (**Supplementary Figure 4A-B**). Finally, when examining differences in organoid morphology regardless of hydrogel or media conditioning (**Figure 2F**), the organoids clustered clearly into three distinct morphological categories: cystic, solid, and mesenchymal.

Among the morphological descriptors used by the classifier (**Table 3**), circularity, convexity, sphericity, geodesic elongation, and lumen radius were the most significant features for classifying organoid morphologies based on their κ-statistic value. We analyzed these features, as well as the volume, as a factor of organoid morphology. Cystic forms were consistently larger than both solid and invasive types in non-TGFβ-treated hydrogels and outsized all organoids found in TGFβ-treated conditions (**Figure 2G**). Regarding circularity (**Figure 2H**), invasive phenotypes predictably exhibited lower circularity than solid or cystic organoids, regardless of hydrogel type. Notably, Matrigel-derived cystic and solid organoids displayed significantly higher circularity compared to those in CM and C hydrogels. This trend also extended to convexity and sphericity and inverted to the geodesic elongation (**Supplementary Figure 4C-E**). Finally, lumen radius values were aligned consistently with organoid volume (**Supplementary Figure 4F**).

### EMT status in PDAC organoids is influenced by ECM composition and TGFβ

Next, we explored if the architecture and composition of the 3D matrix affects the transcriptional state of a custom EMT gene signature: E-cadherin (*CDH1*), N-cadherin (*CDH2*), Vimentin (*VIM*), and *ZEB1* (Palamaris et al., 2021). A two-way MANOVA test of our qRT-PCR results revealed significant main effects on RNA expression levels driven both by matrix composition (ηp^2^ = 0.623, p < 0.001) and the presence of TGFβ in the media (ηp^2^ = 0.883, p < 0.001). Additionally, a significant yet moderate interaction was observed between these two factors (ηp^2^ = 0.383, p < 0.001). Subsequent univariate ANOVA analyses identified *CDH1* (ηp^2^ = 0.717) and *ZEB1* (ηp^2^ = 0.747) as the genes more significantly influenced by matrix composition, with hydrogel C causing the highest effect on gene expression. Regarding media conditioning, *ZEB1* (ηp^2^ = 0.768) and *CDH1* (ηp^2^ = 0.672) are also the most affected genes, with TGFβ exerting the highest impact on the EMT signature compared to the regular media.

Tukey’s corrected post-hoc analysis revealed significant down-regulation of the epithelial gene *CDH1* and concurrent over-expression of mesenchymal markers (*CDH2*, *VIM*, and *ZEB1*) when transitioning from 2D to 3D cultures, particularly within the C hydrogel or under TGFβ conditioning (**Figure 3A-D**). When looking at 3D scaffolds only, no significant differences in *CDH1* expression were observed among hydrogels (**Figure 3A**). Contrarily, *CDH2*, *VIM*, and *ZEB1* expression levels were significantly up-regulated in C hydrogels compared to their M and CM counterparts (**Figure 3B-D**). Finally, the presence of TGFβ caused consistent increases in the expression of all the genes analyzed between organoids cultured within the same type of hydrogel. These results are illustrated in the hierarchical heat-map (**Figure 3E**), which shows two primary gene clusters (*CDH1* and *CDH2*/*VIM*/*ZEB1*) and highlights how the down-regulation of *CDH1* and the up-regulation of *CDH2*/*VIM*/*ZEB1* are pivotal for EMT transition. The heat map further highlights the impact of TGFβ on the gene signature, except for hydrogel C, where EMT is already induced by the gel’s composition.

**Figure 3.**
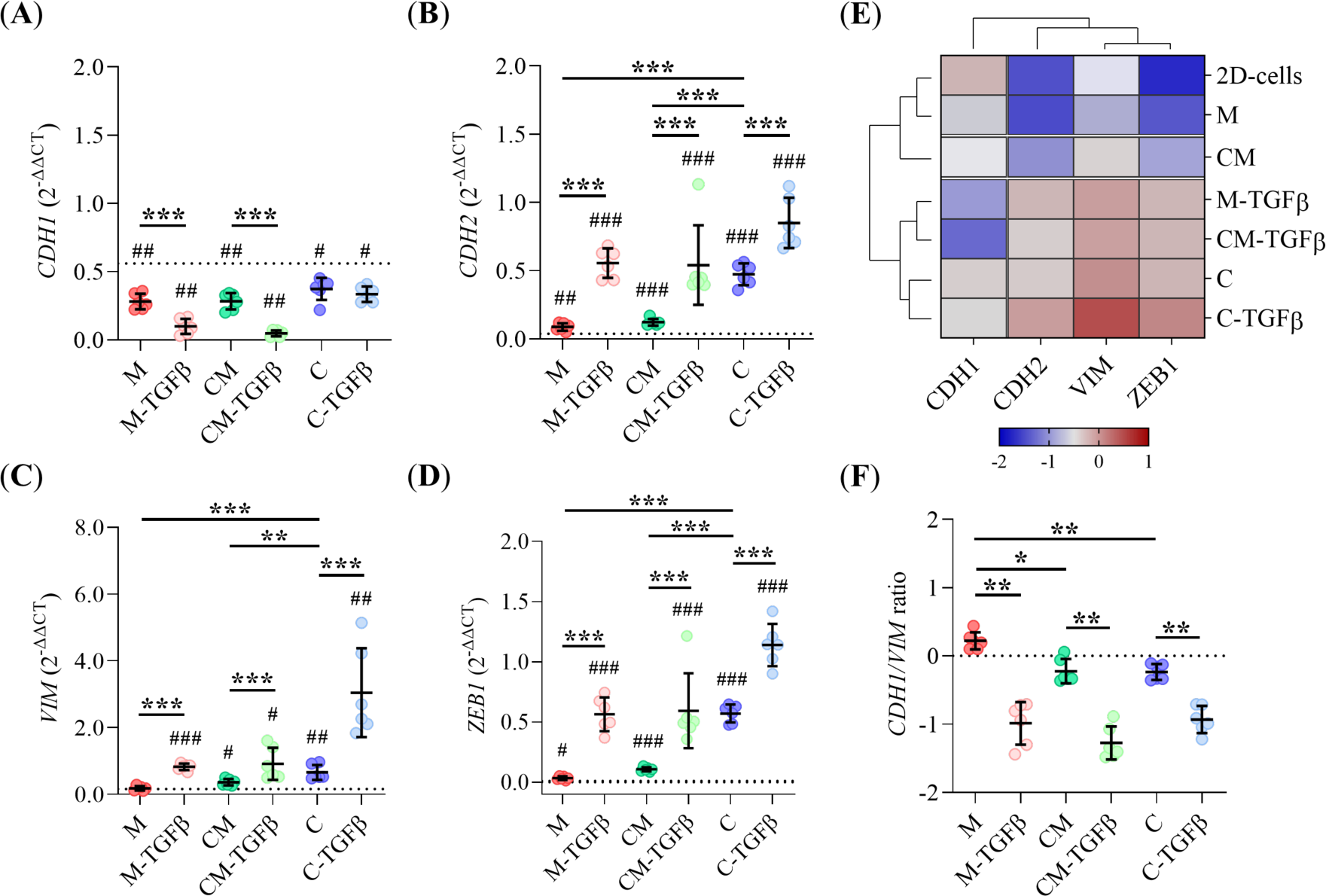
qRT-PCR analysis of EMT-associated genes. (**A**-**D**), Relative mRNA expression levels of *CDH1*, *CDH2*, *VIM*, and *ZEB1*, respectively, in PDAC organoids cultured in M, CM, and C hydrogels under regular media or TFGβ conditioning. mRNA levels were normalized to GAPDH and calculated using the 2^−ΔΔCT^ method. The dotted line represents mRNA expression levels in 2D-plated cells. (**E**) Hierarchical heat-map showing the expression profiles of EMT markers in PDAC organoids, as extracted from panels A to D. (**F**), Quantification of *CDH1*/*VIM* ratio. The dotted line represents the threshold distinguishing epithelial from mesenchymal status. *** Indicates highly statistically significant differences (p<0.001). ** Indicates moderately statistically significant differences (p<0.01). * Indicates statistically significant differences (p<0.05). (#) Refer to the differences compared to 2D-plated cells.

The *CDH1*/*VIM* ratio (El Amrani et al., 2019a) confirms that organoids in CM and C hydrogels have a potential mesenchymal phenotype (*CDH1*/*VIM* ratio < 0), while those in the M hydrogel retained an epithelial one (*CDH1*/*VIM* ratio > 0) (**Figure 3F**). In alignment with these findings, the heat-map also reveals that the gene signature of the 2D cultures closely resembles that of organoids grown in Matrigel (**Figure 3E**). In contrast, CM organoids fall between M and C organoids, which exhibit marked EMT reprogramming. Finally, the presence of TGFβ consistently shifts the genetic signature of all organoids toward a more pronounced EMT profile.

Based on these results, and to streamline our data interpretation in subsequent assays, we renamed cystic organoids as epithelial-like organoids (ELOs) and grouped solid and invasive organoids into a single category labeled mesenchymal-like organoids (MLOs). This classification is supported by the correlation between organoid morphology and their genetic signatures: M-organoids, which predominantly exhibit cystic morphologies, displayed an epithelial-like signature closely resembling that of 2D cultures, while conditions containing mostly solid and invasive morphologies, including those treated with TGFβ, clustered under a unified mesenchymal-like signature.

### Invasive organoid morphologies trigger enhanced ECM remodeling

To study the ECM remodeling caused by PDAC organoids, we analyzed collagen-I fiber deposition, cross-linking, and alignment within the hydrogels. Representative SHG images of the collagen-I mesh remodeling are shown in **Figure 4A-B**. A schematic representation of the image analysis procedure used is depicted in **Figure 4C**. As shown by our results, collagen-I densification by both ELOs and MLOs, either conditioned with TGFβ or not, was evident in all hydrogel types (**Figure 4D**). However, collagen-I densification levels were markedly higher around organoids grown in mixed-composition hydrogels (CM-ELOs and CM-MLOs) compared to those grown in collagen-only (C-ELOs and C-MLOs). Within CM hydrogels, CM-ELOs, mostly associated with cystic forms, displayed higher densification compared to their mesenchymal-like counterparts (CM-MLOs). Matrix alignment, measured through the anisotropy index (α), was detected in MLOs but not ELOs, irrespective of the hydrogel (**Figure 4E**). Among MLOs, CM-MLOs produced enhanced collagen-I fiber alignment compared to C-MLOs. Finally, TGFβ conditioning significantly increased collagen-I fiber alignment in PDAC organoids within CM hydrogels.

**Figure 4.**
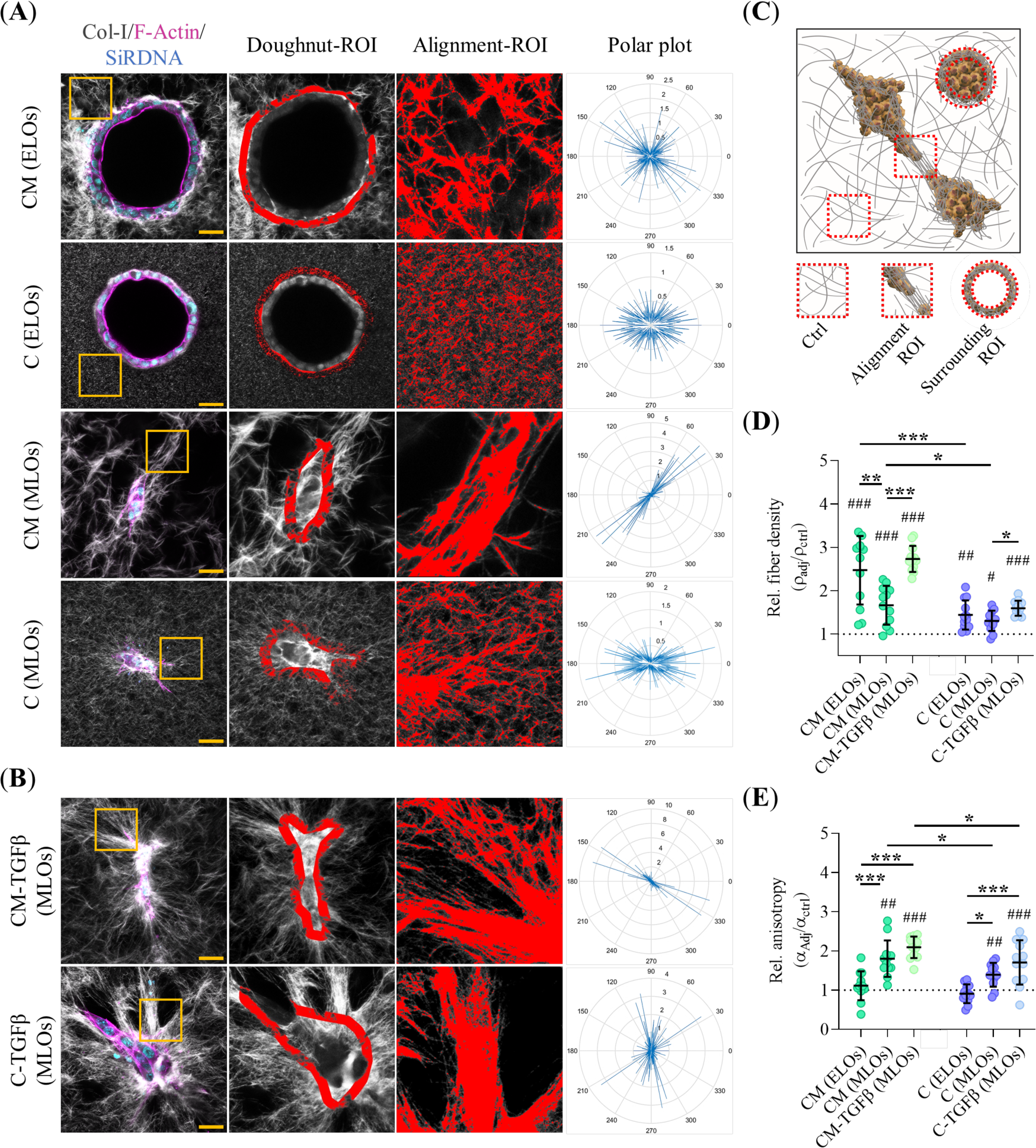
Collagen-I Remodeling in PDAC Organoids. (**A**-**B**), Z-stack projections of representative 3D second harmonic MP-CLSM images of collagen-I remodeling in the different PDAC organoid classes cultured in CM and C hydrogels under regular or TFGβ conditioning, respectively. Staining: F-Actin (magenta) and nuclei (SiRDNA, cyan). The doughnut ROI illustrates fiber segmentation surrounding the organoids (7-micron band). Yellow square insets show representative areas with fiber remodeling. The alignment ROI and the polar plots highlight fiber alignment extracted from the yellow square inset. Scale bars: 25 µm. (**C**), Schematic representation of organoids (orange), fibers (gray), and the control, alignment, and surrounding ROIs used for quantification. (**D**-**E**), Quantification of fiber compaction and alignment (anisotropy, α), respectively, extracted from panels A and B. The dotted line represents fiber remodeling in the control ROI. *** Indicates highly statistically significant differences (p<0.001). ** Indicates moderately statistically significant differences (p<0.01). * Indicates statistically significant differences (p<0.05). (#) Refer to the differences compared to 2D-plated cells.

### Hydrogel composition and TFGβ influence EMT marker location on PDAC organoids

To better understand the effect of the hydrogel on the EMT transition of PDAC organoids, we analyzed the subcellular localization of *CDH1*, *VIM*, and the cell-cell junction protein β-catenin (*CTNNB1*). **Figure 5A-B** illustrates the localization of these proteins in the most frequent organoid types found within each hydrogel type, both with and without TGFβ conditioning. The quantitative analysis of the junction-to-cytoplasm ratio of *CDH1* revealed a marked shift toward the cytoplasm in MLOs compared to ELOs, regardless of matrix composition or media conditioning (**Figure 5C**). Interestingly, no significant differences in *CDH1* distribution were observed within the same organoid type across different hydrogels or after TGFβ conditioning. This is further illustrated in the CLSM images, which show a uniform distribution of *CDH1* along cell-cell junctions in ELOs, while MLOs exhibit a patchy distribution of *CDH1* in the cytoplasm (**Figure 5A-B** and **Supplementary Figure 5**). Besides, the junction-to-nucleus ratio of *CTNNB1* mirrored the pattern observed for *CDH1*, with MLOs exhibiting a marked shift of *CTNNB1* from the membrane to the nucleus complex compared to ELOs, regardless of matrix composition or media conditions (**Figure 5D**). Nevertheless, *CTNNB1* delocalization was significantly stronger in C-MLOs than in their CM counterparts. Additionally, TGFβ conditioning further amplified *CTNNB1* translocation in MLOs relative to their non-conditioned counterparts.

**Figure 5.**
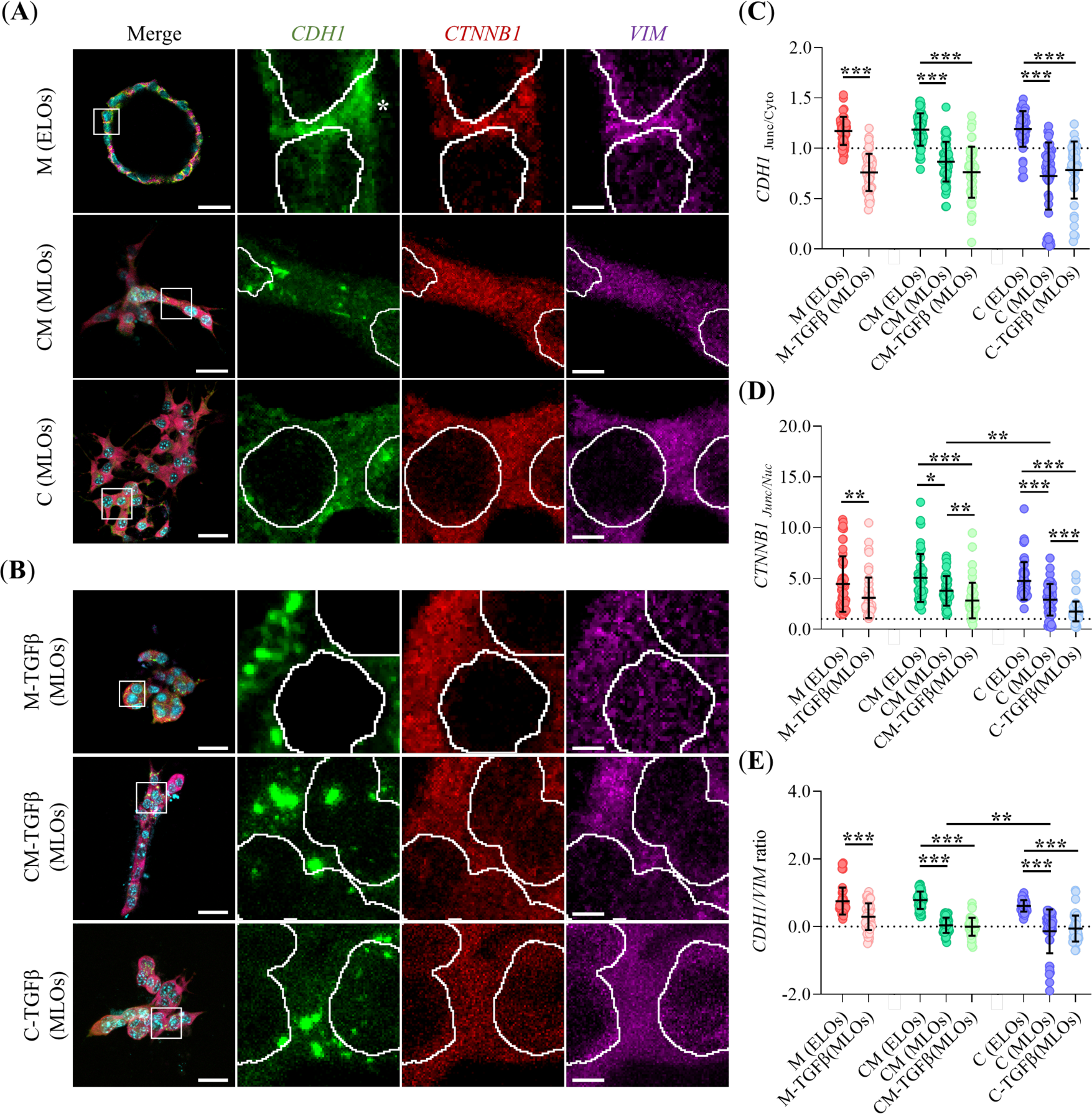
EMT marker distribution analysis. (**A-B**), Z-stack projections of representative CLSM images in the different PDAC organoid classes cultured in CM and C hydrogels under regular or TFGβ conditioning, respectively. Staining: E-cadherin (*CDH1*, green), β-catenin (*CTNNB1*, red), vimentin (*VIM*, magenta), and nuclei (SiRDNA, cyan). Scale bars 30 µm. White square insets provide a zoomed-in view of cell-cell junctions. White asterisks indicate the location of *CDH1* at the cell-cell junctions. Scale bars: 3 µm. (**C-D**), Quantification of EMT marker distribution extracted from the images in panels A and B, including the *CDH1* Junctional/Cytoplasm ratio and the *CTNNB1* Junctional/Nuclei ratio, respectively. The dotted line indicates a ratio of 1. (**E**), Quantification of the *CDH1*/*VIM* ratio. The dotted line represents the threshold distinguishing epithelial from mesenchymal status. *** Indicates highly statistically significant differences (p<0.001). ** Indicates moderately statistically significant differences (p<0.01). * Indicates statistically significant differences (p<0.05).

Finally, the *CDH1*/*VIM* ratio (**Figure 5E**) revealed that both CM-MLOs and C-MLOs, treated or not with TGFβ, displayed a potential mesenchymal phenotype (*CDH1*/*VIM* ratio ≤ 0). In contrast, ELOs consistently retained epithelial identity (*CDH1*/*VIM* ratio > 0), regardless of matrix composition. Interestingly, TGFβ conditioning did not induce a mesenchymal shift in organoids grown within M hydrogel (*CDH1*/*VIM* ratio > 0). Overall, while *CDH1* qRT-PCR levels remained consistent across all tested hydrogels, its subcellular localization varied significantly. Notably, *CDH1* delocalization in collagen-containing hydrogels, along with *CTNNB1* translocation, highlights the influence of matrix composition on EMT progression.

### ECM stiffening rewires the metabolic and mitochondrial remodeling in PDAC organoids

To investigate the impact of the biomechanical properties of the ECM on the metabolism of PDAC organoids, we monitored their cellular oxygen consumption rate (OCR) and their lactate production through extracellular acidification rate (ECAR). Our results show that organoids grown in CM hydrogels exhibit heightened basal and maximal respiration rates, along with enhanced SRC, compared to their counterparts grown in M and C hydrogels (**Figure 6A, C**). Moreover, CM hydrogels showed higher ECAR rates, indicating increased glycolytic activity across basal, maximal, and GC rates compared to M hydrogel (**Figure 6B, D**). No ECAR differences were detected between the CM and C hydrogels, except for the basal glycolysis rate (**Figure 6B, D**). Furthermore, Mito- and Glyco-ATP production rates revealed that PDAC organoids primarily rely on oxidative phosphorylation (OXPHOS) for energy, regardless of the ECM composition (**Figure 6E**). However, our results confirmed that PDAC organoids also utilize aerobic glycolysis as an additional energy source. Interestingly, Glyco-ATP production was significantly higher in CM hydrogels compared to M and C hydrogels (**Figure 6E**). Altogether, the metabolic profile of the PDAC organoids, based on the basal OCR-ECAR ratio, reveals short energetic demands for M and C organoids and a high energy-consuming phenotype for CM organoids (**Figure 6F**).

**Figure 6.**
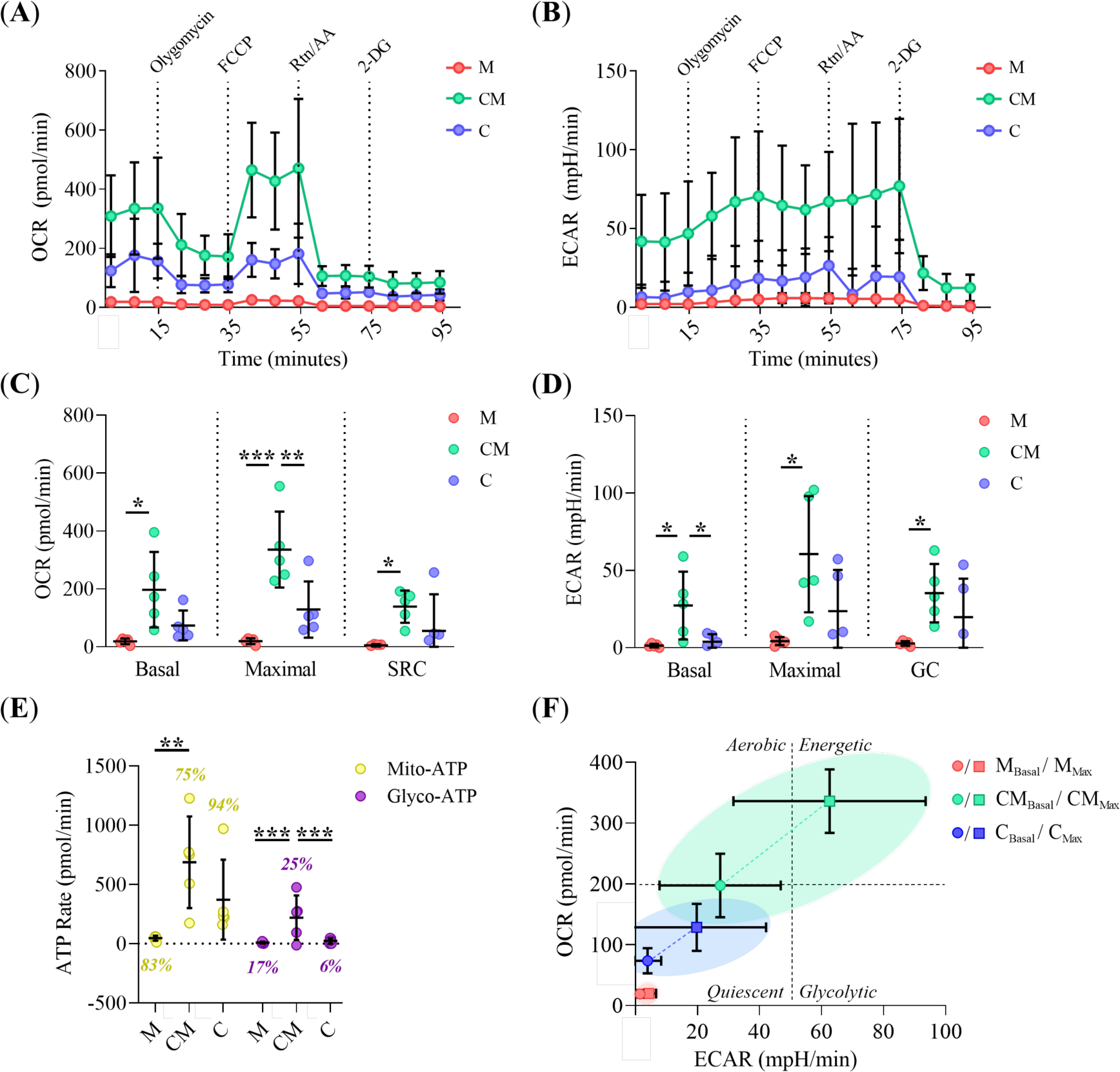
Bioenergetic analysis of PDAC Organoids. (**A, B**), Representative OCR (pmol/min) and ECAR (mpH/min) curves over time in PDAC organoids cultured in M, CM, and C hydrogels. (**C**), Basal and maximal respiration rates, as well as SRC (pmol/min), extracted from panel A. (**D**), Basal and maximal proton flux, as well as GC rate (mpH/min), extracted from panel B. (**E**), ATP production rate (pmol/min), showing contributions from mito-ATP (yellow) and glycol-ATP (magenta). Percentages represent the ATP generated by each pathway relative to total ATP production. (**F**), Energetic map depicts basal (circle) and maximal (square) OCR and ECAR levels in PDAC organoids grown in M, CM, and C hydrogels. The four quadrants represent metabolic states: “Quiescent,” “Aerobic,” “Energetic,” or “Glycolytic.” Colored ellipses illustrate the metabolic space occupied by organoids within each hydrogel condition. *** Indicates highly statistically significant differences (p<0.001). ** Indicates moderately statistically significant differences (p<0.01). * Indicates statistically significant differences (p<0.05).

To better understand the metabolic changes observed across the different hydrogels, we analyzed mitochondrial morphology and its membrane potential (Δψm) in PDAC organoids. CLSM images (**Figure 7A**) revealed a significant increase in mitochondrial mass in organoids cultured in CM and C hydrogels compared to M organoids (**Figure 7B**). In addition, CM- and C-grown organoids exhibited enlarged and hyper-fused mitochondria, based on the level of branches, in comparison to M-grown organoids (**Figure 7C-E**). No significant differences in mitochondrial sphericity were observed between the tested hydrogels (**Figure 7D**). Conversely, CM and C mitochondria displayed enhanced Δψm (**Figure 7F-G**) compared to M organoids. Remarkably, 2D cultures exhibited lower mitochondrial mass, mean volume, fusion level, and Δψm than PDAC organoids (**Figure 7B-G**), regardless of the matrix composition. Moreover, compared to 3D matrices, 2D environments promote mitochondrial sphericity, indicating dysfunctional mitochondria (**Figure 7D**). All these findings suggest that 3D environments boost mitochondrial mass and metabolic rewiring, especially within CM hydrogel. More importantly, CM organoids adopt a highly energetic phenotype sustained by robust oxidative phosphorylation and aerobic glycolysis, consistent with the Warburg effect.

**Figure 7.**
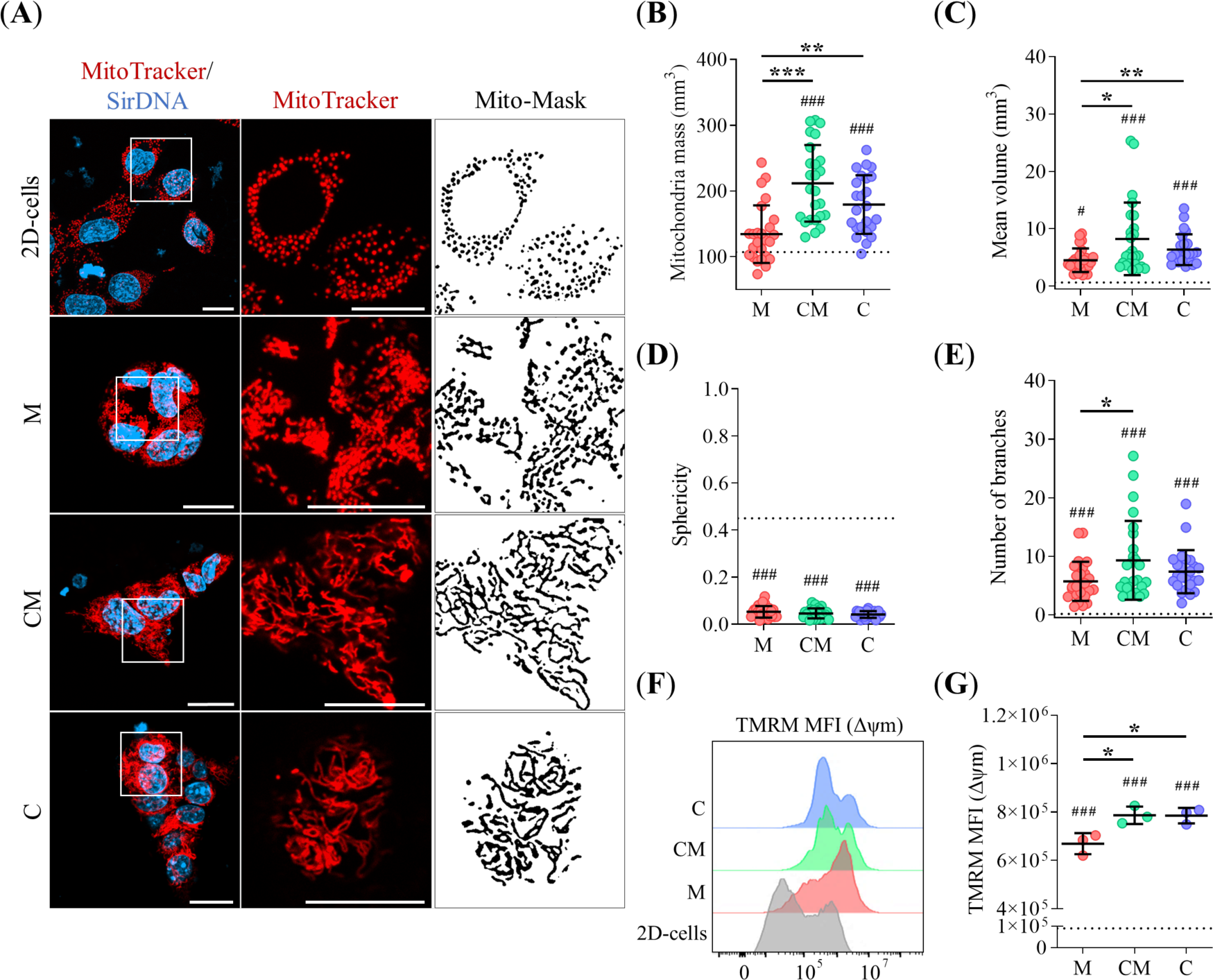
Mitochondrial analysis in PDAC organoids. (**A**), Z-stack projections of representative CLSM images of PDAC organoids cultured in CM and C hydrogels alongside 2D-plated cells under standard media conditions. Staining: mitochondria (MitoTracker, red) and nuclei (SiRDNA, cyan). White square insets display magnified regions highlighting mitochondrial morphology. Scale bars: 30µm. The mask column shows the mitochondria segmentation. Scale bars: 25 µm. (**B-E**), Quantification of mitochondrial morphological descriptors, including mitochondrial mass (mm^3^), mean volume (mm^3^), sphericity, and number of branches, respectively, extracted from panel A. The dotted line indicates levels in 2D-plated cells. (**F-G**), Representative FACS histograms and MFI values for TMRM staining (Δψm) in PDAC organoids cultured in M, CM, and C hydrogels alongside 2D-plated cells. The dotted line indicates levels in 2D-plated cells. *** Indicates highly statistically significant differences (p<0.001). ** Indicates moderately statistically significant differences (p<0.01). * Indicates statistically significant differences (p<0.05). (#) Refer to the differences compared to 2D-plated cells.

### 3D-environment confers adaptive Gem-resistance in PDAC organoids

To explore the impact of our hydrogels on PDAC organoid drug resistance, we performed *in vitro* live/dead assays under Gem conditioning. Prior to this, we established a GI50 dose-response curve using Matrigel, which was expected to have the highest chemo-sensitivity based on its EMT phenotype (**Figure 8A-B**). Representative images of the assay for all experimental settings are shown in **Figure 8C-D**. Our findings indicate that PDAC organoids cultured within CM and C hydrogels exhibited approximately 40% higher resistance to Gem compared to those grown on M hydrogels (**Figure 8E**). Notably, the live/dead ratio revealed no significant differences in Gem sensitivity between CM and C organoids, with approximately 10% mortality, regardless of the media used. TGFβ conditioning significantly increased the viability of M organoids, bringing their resistance levels close to those observed in CM and C hydrogels. Interestingly, no significant differences in Gem sensitivity were observed between 2D cultures and organoids cultured within the M hydrogel.

**Figure 8.**
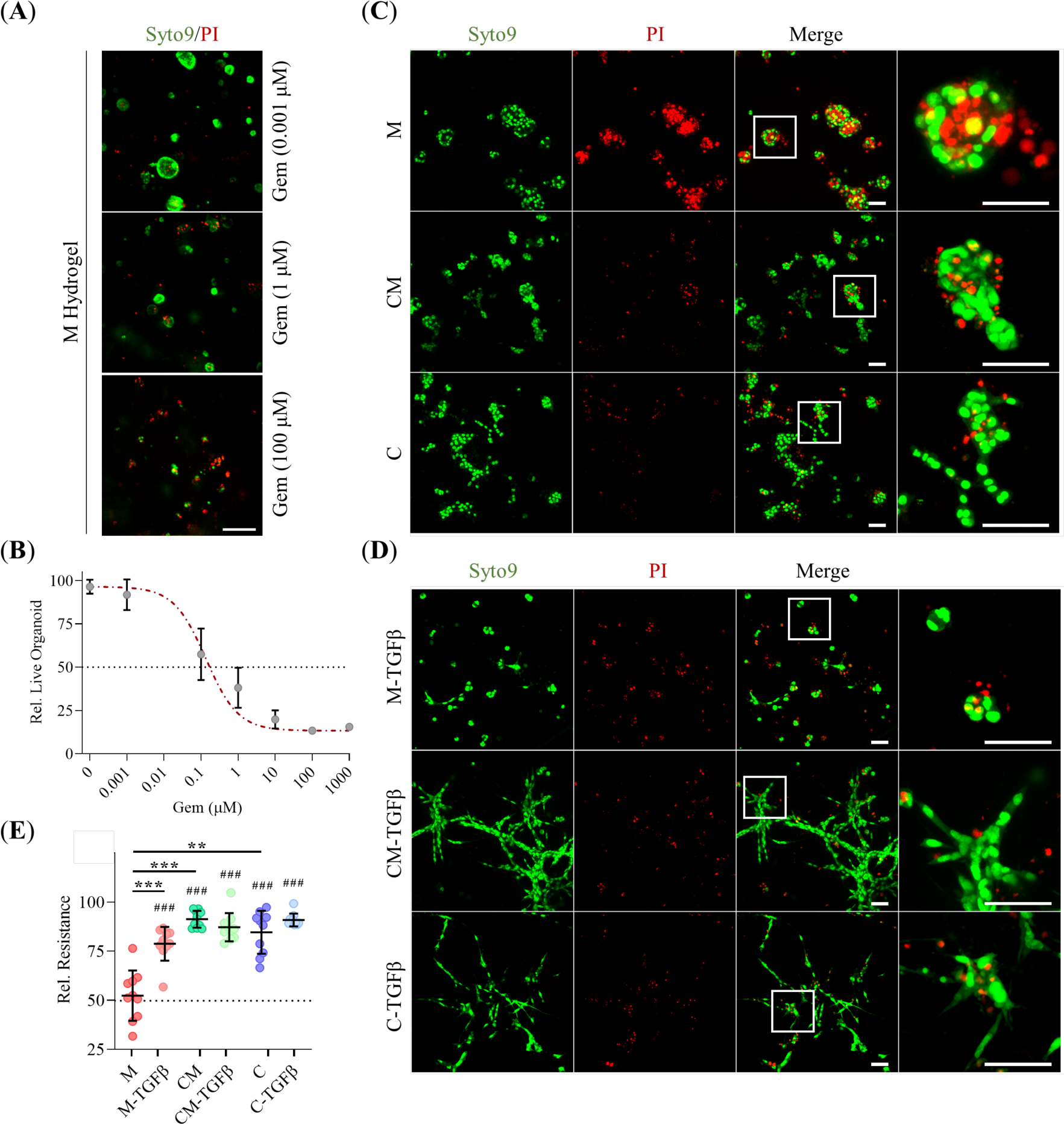
*In vitro* Gem resistance in PDAC organoids: (**A**), Z-stack projections of representative CLSM images of PDAC organoids cultured in M hydrogels treated with increasing concentrations of Gem (0.001, 1, and 100 μM). Staining: live nuclei (Syto9, green) and necrotic nuclei (PI, red). Scale bars: 150 µm. (**B**), IG50 dose-response curve extracted from the images in panel A. The dotted line indicates the IG50 value. (**C-D**), Z-stack projections of representative CLSM images of PDAC organoids cultured in M, CM, and C hydrogels, under regular or TFGβ conditioning, respectively, in the presence of 0.1 µM Gem. Staining: live nuclei (Syto9, green) and necrotic nuclei (PI, red). High-magnification white insets provide detailed views of organoid status under each condition. Scale bars: 30 µm. (**E**), Quantification of organoid chemoresistance to Gem under regular or TFGβ conditioning, respectively. The dotted line represents sensitivity levels in 2D-plated cells. *** Indicates highly statistically significant differences (p<0.001). ** Indicates moderately statistically significant differences (p<0.01). * Indicates statistically significant differences (p<0.05). (#) Refer to the differences compared to 2D-plated cells.

### ECM composition influences tumor growth and Gem-resistance in PDAC organoid-derived cancer-bearing mice

Finally, we examined how the matrix biomechanical properties influence tumor growth, neovascularization, and Gem response in immunocompetent mice-bearing tumors. To this end, PDAC organoids grown in M, CM, and C hydrogels were injected into syngeneic C57BL/6 mice (n = 9/group). Metronomic Gem treatment was initiated when the tumor volume reached 50 mm^3^ (day 16) and was monitored by echography on day 28 (**Figure 9A**). At this point, PDAC tumors derived from organoids embedded in CM and C hydrogels exhibited significantly larger normalized volumes than those grown in M hydrogels (**Figure 9B-D**). In contrast, Gem treatment substantially reduced tumor volume in tumors derived from organoids cultured within M and CM hydrogels (∼80% and ∼70%, respectively) compared to untreated tumors. No significant differences in tumor volume were found between the C and C-Gem groups (**Figure 9C-D**). These results were confirmed by tumor images harvested on day 28 (**Figure 9E**). Besides, CM- and C-derived tumors exhibited increased peri- and intra-tumoral vasculature compared to M-derived tumors (**Figure 9B-F**), while metronomic Gem enhanced tumor vascular density across all hydrogel types compared to untreated tumors (**Figure 9C-F**). To further evaluate Gem response, we calculated the tumor growth inhibition (TGI) index, which revealed significantly higher growth inhibition in Gem-treated tumors derived from M organoids compared to those grown in CM and C hydrogels (**Figure 9G**). No significant differences in Gem sensitivity were observed between CM-Gem and C-Gem groups, with C-derived tumors displaying particular chemoresistance to Gem. These results suggest that CM hydrogels provide an adequate scaffold for PDAC tumor development and that the presence of collagen-I boosts tumor chemoresistance in our murine model.

**Figure 9.**
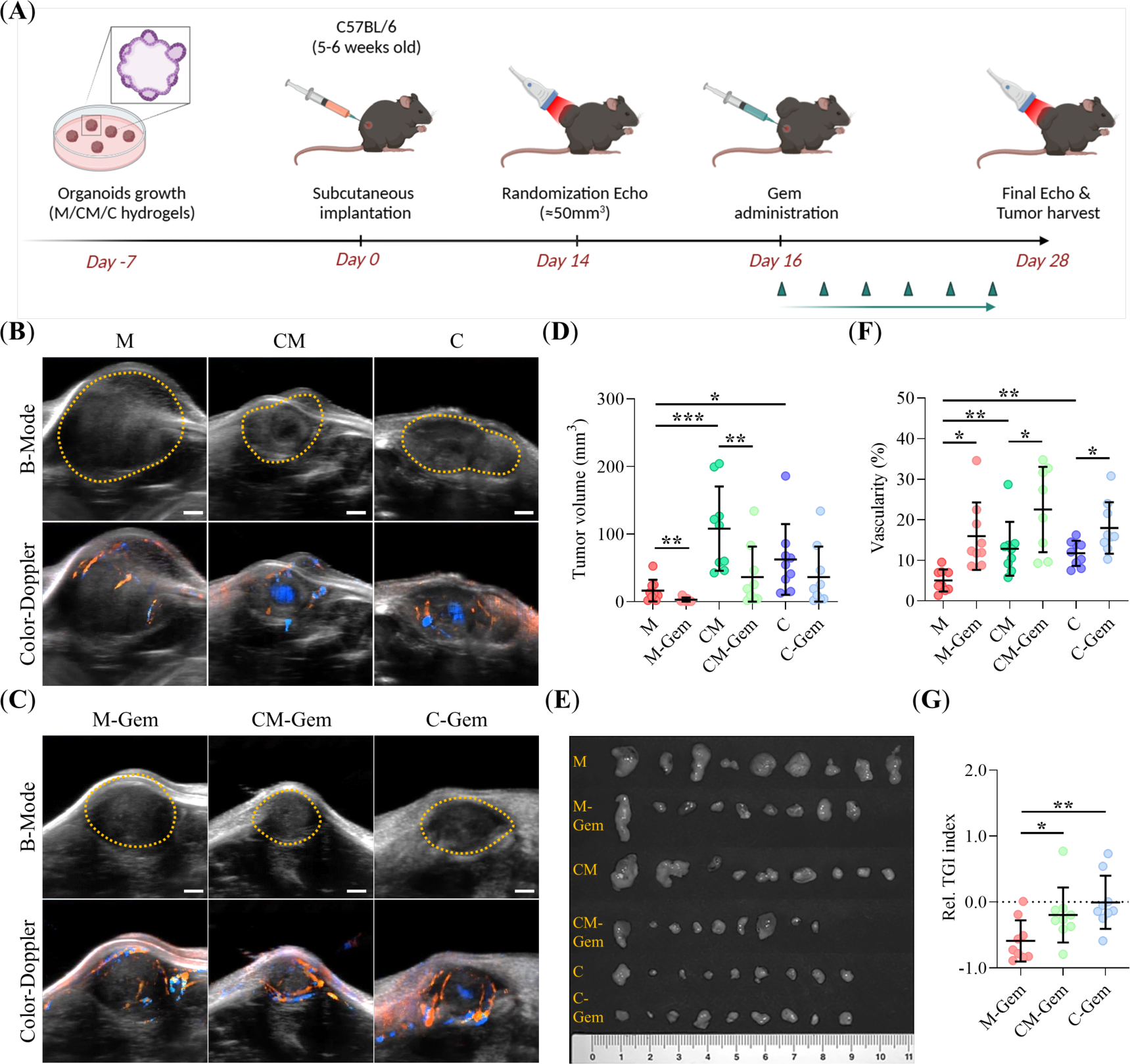
*In vivo* Gem resistance in a KPC mice model. (**A**), Schematic of the experimental design. Metronomic Gem (100 mg/kg) was administered intraperitoneally on days 16, 18, 20, 22, 24, and 26 (green triangles). (**B-C**), Representative 3D-rendered views of tumors 28 days after organoid injection, comparing groups with and without Gem treatment. B-mode images show anatomical structure alongside the surrounding and intra-tumoral vasculature (Color Doppler-mode, red and blue). The dotted yellow line indicates the tumor margins. Scale bars: 1 mm. (**D-E**), Quantitative analysis of tumor volume and vasculature, respectively, extracted from panel B and C. (**F**), Photographs of excised tumors harvested on day 28. (**G**), Quantification of the relative tumor growth inhibition index (TGI) in the Gem-treated groups. The dotted line indicates the threshold separating tumor regression (TGI < 0) and tumor progression (TGI > 0). *** Indicates highly statistically significant differences (p<0.001). ** Indicates moderately statistically significant differences (p<0.01). * Indicates statistically significant differences (p<0.05).

## Discussion

The biomechanical properties of the ECM play a critical role in tumor invasion, angiogenesis, metastasis, and drug resistance (Liu et al., 2023; Popova & Jücker, 2022; Yuan et al., 2023). Despite this, most cancer studies are conducted using conventional 2D cultures or in suspension-based 3D models, such as cancer spheroids (Cardoso et al., 2023). Recently, 3D tumor organoid models have gained popularity because of their tumor resemblance and translational potential. However, these organoid models often use scaffolds with unreported or poorly characterized biomechanical properties (Barbosa et al., 2021; Hoarau-Véchot et al., 2018), severely limiting their interpretability. In this work, we aim to address these limitations by providing a comprehensive analysis of how the scaffold biomechanics influence the morphology, phenotype, metabolism, and drug resistance of murine PDAC organoids. To this end, we used three hydrogel types: Matrigel-only (M), representing a biochemically rich but mechanically soft environment; collagen-only (C), representing a structured, rigid, bio-functional but biochemically poor environment; and a mixture of collagen-Matrigel (CM), representing an environment with both structural and biochemical complexity.

To accurately interpret the effect of these matrices on PDAC organoid biology, we first measured the mechanical properties of the hydrogels. Collagen-only hydrogels exhibit high G’, typical of rigid hydrogels with enhanced elastic behavior. These hydrogels also display a high G’’, indicating a high viscous component. This mechanical profile has been reported to promote cell migration by enabling cells to generate tensions at rapid time scales and promoting ECM remodeling (Krajina et al., 2021). Consistent with this, our C hydrogels displayed the highest invasive capacity of EOSs among the three hydrogels tested. This enhanced 3D migration in increasingly rigid scaffolds, driven by collagen-I concentration, has been well-documented to promote collective cell migration and mechanical stability, facilitating organoid development in various cancer types, including liver (M. Kim et al., 2023), colon (Jee et al., 2019), breast (Bhattacharya et al., 2023) or glioblastoma (Sun et al., 2024).

Matrigel contributes to the rigidity of the hydrogels but less so than collagen at an equal concentration. Moreover, the higher the concentration of Matrigel, the lower the relative rigidity and viscosity of the hydrogel (Borries et al., 2020). This is consistent with the lowest migration dynamics of the EOSs found in M hydrogels and explains the lower migration dynamics seen in mixed CM hydrogels compared to collagen-only C hydrogels. However, the reported reduction found is smaller than expected. These findings can be attributed not only to the larger pore size of CM hydrogels compared to C but also to the mechanobiological changes introduced by Matrigel, which affect the way cells migrate. Indeed, we have previously demonstrated (Anguiano et al., 2020) that low concentrations of Matrigel enhance H1299 lung cancer cell migration by stimulating β1 integrin expression and the formation of focal adhesions, which enhance cell traction. However, excessive Matrigel leads to too-large adhesion formation, which favors attachment over traction and ultimately slows migration rates (Anguiano et al., 2020; Poincloux et al., 2011). TFGβ conditioning enhances EOS migration speed and protrusion formation in all scaffolds, particularly in collagen-containing (C, CM) scaffolds compared to Matrigel-only ones.

Our results of EOS migration align with the morphologies observed in the mature organoids. In M scaffolds, organoids are mostly cystic, while those in CM or C hydrogels exhibit invasive or solid forms. This indicates that the more static EOSs, predominantly found in M hydrogels, self-organize into large cystic forms, while highly motile EOSs in CM and C hydrogels form smaller organoids with either solid or invasive morphologies. The TFGβ-induced trend towards invasiveness seen in EOSs also correlates with the invasive morphologies seen in mature organoids, as solid and invasive forms are the most frequent, irrespective of the matrix composition. Finally, this enhanced pro-migratory phenotype is consistent with the reduced number and the smaller size of the organoids grown with TFGβ.

We next examined EMT marker expression in mature organoids as a function of hydrogel composition. EMT involves the replacement of epithelial cytokeratin with mesenchymal vimentin and is also associated with the depletion of *CDH1* at the cell junctions and the activation of *CDH2* (Lamouille et al., 2014). EMT is orchestrated by transcription factors like *SLUG*, *SNAIL*, *TWIST*, and *ZEB1*, which disrupt apical-basal polarity, induce metabolic rewiring (Buckley & St Johnston, 2022), and contribute to tumor aggressiveness and drug resistance (Bakir et al., 2020; De Las Rivas et al., 2021; Kaemmerer et al., 2021). Our qRT-PCR analysis confirmed significant changes in the expression of *CDH2*, *ZEB1* and *VIM* in collagen-I scaffolds. This is consistent with recent studies showing that rigid matrices up-regulate *ZEB1*, and *VIM*, contrarily to softer matrices like Matrigel or in 2D cultures (Liu et al., 2024). While *CDH1* expression decreased with dimensionality, no significant differences were observed across 3D hydrogel compositions. This contrasts with recent studies linking PDAC tumor rigidity to *CDH1* suppression and poor clinical outcomes (Dardare et al., 2021; Rice et al., 2017; Sommariva & Gagliano, 2020). Although CDH1 mRNA levels remain similar across hydrogel types, our immunofluorescence analysis revealed *CDH1* delocalization from the membrane to cytoplasmic patches, particularly in MLOs, irrespective of the matrix composition or medium. This supports the findings by Aiello et al. (Aiello et al., 2016), who identified *CDH1* loss in PDAC tumors as membrane delocalization rather than transcriptional down-regulation. Similarly, it has been reported that pancreatic carcinomas retain *CDH1* expression despite exhibiting high invasiveness and metastatic potential (Sommariva & Gagliano, 2020). We also observed *CTNNB1* delocalization from the membrane to the cytoplasm or nucleus, particularly in MLOs, irrespective of the matrix composition or medium. Although complete nuclear translocation was not observed, cytoplasmic accumulation was evident as an early event in PDAC tumorigenesis and is a well-established hallmark of EMT across various cancer types (Arensman et al., 2014)(W. K. Kim et al., 2019). These findings align with observations by Zhang et al., who reported similar *CTNNB1* patterns in PanIN lesions in KPC mice (Zhang et al., 2013). In summary, organoids grown in collagen-only hydrogels exhibit a pure mesenchymal phenotype, while those in mixed-composition (CM) hydrogels displayed an intermediate phenotype, and Matrigel-only hydrogels retained an epithelial phenotype, resembling cells in 2D culture. *CDH1*/*VIM* ratios, as assessed by both qRT-PCR and immunofluorescence, further validate these findings (El Amrani et al., 2019a). Consequently, we classified cystic organoids as epithelial-like organoids (ELOs) and grouped solid and invasive organoids under the mesenchymal-like organoids (MLOs) category. TGFβ-conditioned hydrogels, as expected (Rajagopal et al., 2021), consistently induced a pure mesenchymal phenotype, marked by strong *ZEB1* and *VIM* over-expression and concomitant *CDH1* loss. Both *CDH1* and *CTNNB1* were also delocalized from the membrane under these conditions, underscoring the role of TGFβ in EMT in PDAC tumors.

ECM remodeling is crucial in PDAC progression (Piersma et al., 2020). Our analysis showed collagen-I densification around organoids across all hydrogel types, with the strongest effects in mixed-composition hydrogels (CM-ELOs and CM-MLOs). However, collagen-I alignment occurred exclusively in MLOs, regardless of hydrogel composition, with the most pronounced effect in CM hydrogels and under TGFβ conditioning. These findings align with reports that identified *ZEB1* up-regulation as a driver of collagen-I remodeling through LOX-dependent activation (Peng et al., 2017). Indeed, in PDAC tumors, stromal populations accumulate extensively fibrous ECM components, promoting EMT and further ECM remodeling (Scott et al., 2019). This heterogeneous deposition and linearization of fibers in the TME promotes cell invasiveness, as seen during EOS migration in our collagen-based hydrogels (Ahmadzadeh et al., 2017). Our results suggest that mixed-composition (CM) hydrogels best mimic the desmoplastic ECM of PDAC tumors, encompassing both the pre-invasive epithelial and invasive mesenchymal states exacerbated by TGFβ. This supports recent studies reporting that ECM stiffening and/or remodeling activate latent TGFβ signaling, fostering tumor invasiveness, drug resistance, and immune evasion (Bauer et al., 2020).

Further emphasizing the importance of matrix dimensionality and composition, our glucose consumption assays revealed that 3D arrangements elicited enhanced mitochondrial bioenergetics compared to 2D configurations (Tidwell et al., 2022). This effect is particularly true in organoids cultured within hydrogels of mixed composition (CM), which show increased glycolytic activity, consistent with the Warburg effect while maintaining simultaneously elevated OXPHOS levels. This metabolic duality supports PDAC cell dissemination and metastasis (Ali et al., 2024). PDAC cells have been shown to reprogram their metabolism in response to matrix rigidity, amplifying the Warburg effect and tumor malignancy. Specifically, as described in the previous paragraph, EMT-driven metabolic reprogramming in PDAC organoids, mediated by *ZEB1*, was most evident in CM- and C-based hydrogels (Valle et al., 2018). Moreover, under confined, stiff ECM conditions, PDAC cells also exhibited mitochondrial remodeling, including increased mitochondrial mass, fusion, and membrane potential (Urra et al., 2021). Our analysis confirmed these trends in 3D hydrogels, with CM and C hydrogels demonstrating the highest mitochondrial remodeling levels, a hallmark of enhanced metastatic potential in PDAC tumors (Carmona-Carmona et al., 2022). Altogether, these findings suggest that CM hydrogels closely mimic the metabolic and phenotypic behaviors of PDAC tumors, which are characterized by upregulated, aberrant glycolytic flux, even under normoxic conditions. These metabolic adaptations confer both invasive and chemoresistance advantages to PDAC cells, as observed in both patient-derived tissues and mouse models (Zheng et al., 2024).

Finally, we summed up the impact of our matrix composition on PDAC with an analysis of the sensitivity to Gem. Multiple factors contribute to Gem chemoresistance in PDAC tumors, including matrix stiffness and remodeling, EMT, and metabolic rewiring (Gu et al., 2021). Consistent with recent findings, our *in vitro* results showed that collagen-based matrices, with their associated biomechanical traits, correlate with higher chemoresistance to Gem compared to softer matrices like Matrigel or to 2D cultures (Pan et al., 2023). Furthermore, extensive fiber deposition and alignment observed in collagen-based hydrogels further exacerbate Gem resistance by restricting drug diffusion (Gregori et al., 2024; Hsu et al., 2022). Notably, Gem-resistant PDAC cells exhibited higher expression of EMT markers (e.g., *ZEB1*) and reduced levels of *CDH1*, explaining the similar chemosensitivity observed between Matrigel-based hydrogels and 2D cultures (Palamaris et al., 2021; Wang et al., 2019). Indeed, EMT status, often quantified by the *CDH1*/*VIM* ratio, has been proposed as a prognostic factor for PDAC tumors’ response to Gem, correlating with increased organoid viability under TGFβ conditioning (El Amrani et al., 2019a). Similarly, highly bioenergetic profiles, including increased glycolytic flux and mitochondrial mass observed in CM- and C-based hydrogels, are also associated with Gem resistance in PDAC cells (Fu et al., 2021; Masuo et al., 2023). In parallel, *in vivo* results further support this assessment in a more clinically relevant immunocompetent mouse model. Specifically, syngeneic CM-derived subcutaneous tumors closely mimicked the PDAC microenvironment, fostering the highest rates of vascularization and tumor growth. Recent studies have highlighted how ECM stiffness plays a crucial role in TME neovascularization, creating a perivascular niche that facilitates cancer metastasis (Mai et al., 2024). Interestingly, as reported, metronomic Gem administration increased tumor vascular density regardless of matrix composition and significantly reduced metabolic activity, ultimately resulting in slower tumor growth (Yapp et al., 2016).

These results suggest that a dense, fibrotic ECM, primarily composed of collagen, impedes drug diffusion while promoting an aggressive, metabolically active phenotype, which contributes to heightened Gem chemoresistance. Notably, mixed-composition (CM) hydrogels closely replicate the PDAC TME by recreating biomechanical stimuli and bioactive cues that better support the delicate balance between tumor growth and vascular development.

In summary, we have shown that the biomechanical properties of the hydrogel modulate the behavior of PDAC tumor organoids and are critical to creating an environment that replicates the properties of the native tumor environment. In this regard, we have shown that hydrogels of mixed collagen-Matrigel composition provide an environment that mechanically approximates the rigidity and viscosity of the tumor ECM better than pure Matrigel of collagen hydrogels. These biomechanical properties promote a pro-migratory scenario that favors the development of invasive, highly metabolic chemo-resistant tumor phenotypes and the remodeling activity typical of the highly desmoplastic ECM of PDAC tumors. In that regard, CM hydrogels lay between the soft environment of M hydrogels, which many aspects are no different from 2D cell cultures, and the rigid but bare and excessively pro-migratory environment of pure C hydrogels that do not allow the development of realistic tumors.

## Materials and Methods

### 3D printed master mold

A 3D CAD model (**Supplementary Figure 1A**) of the master mold used to fabricate the micro-devices was generated using Inventor 2020 (Autodesk, USA) and 3D-printed in a Form2 printer (FormLabs, USA) using Tough 2000 resin (RS-F2-TO20-01, FormLabs, USA) with an axial resolution of 50 μm (**Supplementary Figure 1B**). The fabricated master mold was washed with 2-propanol in a Form Wash device and post-cured in a Form Cure UV oven at 80°C for 2h.

### Microfabricated device for the growth of organoids

3D organoids were grown in custom-made Polydimethylsiloxane (PDMS) micro-devices containing three 5 mm diameter hydrogel-loading wells enclosed in a 1.2 mL cylindrical media reservoir (see **Supplementary Figure 2A-B**). The reservoirs were fabricated by casting a 10:1 base-to-curing agent ratio PDMS mixture onto the 3D printed master mold described above, followed by degassing for 25 min and polymer curation at 70°C for 2. Subsequently, the cured PDMS was detached from the master mold, and three holes were punched (5 mm) to form the wells for hydrogel loading. PDMS micro-device was then plasma bonded to a glass coverslip (#1 thickness, 35 mm diameter; Menzel-Glaser, Germany) using a Zepto plasma system (Diener Electronic, Germany) at 85 W power and 0.3-0.4 mBar pure O2 pressure for 1 min. **Supplementary Figure 2C** shows one of the fabricated devices. Before any experiment, the devices were UV sterilized for 15 min, and a 0.1 mg/mL Poly-D-lysine coating was applied to enhance hydrogel adherence.

### Hydrogel preparation

Three different hydrogels were prepared using varying concentrations of collagen-I (BD Biosciences, USA) and growth factor-free (GFR) Matrigel (Corning, USA). We refer to them as M (4 mg/mL Matrigel), CM (2 mg/mL collagen-I, 2 mg/mL Matrigel) and C (4 mg/mL collagen-I). To fabricate the hydrogel, a mixture of 10x PBS and sterile deionized water was first prepared, followed by the addition of collagen-I and/or Matrigel at the desired concentration (pH = 7). C and CM hydrogel gelation was performed in two steps: the hydrogels were first incubated at 25°C for 30 min and then at 37°C for 15 min. M hydrogel was gelated in a single incubation step at 37°C for 15 min.

### Hydrogel morphological characterization

The 3D microstructural characterization of hydrogels was conducted through automated analysis of Z-stack images acquired using a Zeiss LSM-880 AxioObserver inverted multiphoton excitation confocal laser scanning microscope (MP-CLSM) equipped with a 63x C-Apochromat objective (1.2 NA, W). The microscope’s Mai-Tai® DeepSee™ T-Sapphire laser was tuned to 790 nm for SHG imaging of the collagen-I fiber network. The fiber signal was filtered through a 380-420 nm band-pass filter before being directed to the Zeiss BiG-2 GaAsP detection module. Z-stacks images comprising a total volume of 135×135×50 µm3 were acquired while keeping the hydrogels in organoid culture media at 37°C and 5% CO2 to simulate culture conditions. The collagen-I network was analyzed using a homemade script for Fiji (Schindelin et al., 2012a). The collagen-I network geometry, including fiber thickness, density, and pore diameter, was analyzed using the pipeline previously described by our group (Anguiano et al., 2020). Ten images were analyzed from three independent experiments.

### Hydrogel rheological characterization

The mechanical properties of the hydrogels were characterized using a stress-controlled DHR-1 rheometer (TA Instruments, USA) with a plate-cone geometry, as previously described (Valero et al., 2018). Briefly, 1.2 mL of the hydrogel prepolymer was placed on the rheometer’s lower plate. The gap between the upper and lower plates was initially set to 0.1 mm and maintained constant throughout the gelation process. Once gelation was complete, the gap was adjusted to 0.052 mm. Then, time sweep experiments were performed for at least 1h, applying to the hydrogels a sinusoidal strain amplitude of 0.3% (linear viscoelastic regime, LVR) at a frequency of 0.1 Hz. The storage (G’) and loss (G”) moduli of the three viscoelastic gels were recorded over time. Three independent experiments were performed per hydrogel type.

### Cell culture

Organoid-generating cells were derived from primary pancreatic tumors in Kras^+/LSL-G12D^; Trp53^+/LSL-R172H^; Pdx1-Cre (KPC) mice on a C57Bl/6 strain background. These organoids produced a range of ductal lesions that closely resemble low- and high-grade human pancreatic intraepithelial neoplasia (PanINs), which can further progress into both primary and metastatic pancreatic ductal adenocarcinoma (PDA) (Boj et al., 2015). Organoid-generating cells were incubated in DMEM medium supplemented with 10% FetalClone III (Cytiva, USA) and 1% Penicillin/Streptomycin (Gibco, Spain) at 37°C and 5% CO_2_. When necessary, organoid-generating cells were pre-conditioned for 48h with regular medium supplemented with 1 ng/mL TGFβ, which was subsequently maintained throughout the entire culture period. A detailed description of the culture media used is available in **Supplementary Table 1**.

### GFP Lentivirus transduction

Stably GFP-expressing organoid-generating cells (PDAC-GFP) were obtained via lentivirus transduction of parental cells. Lentiviruses were produced using X-tremeGENE DNA Transfection Reagent (Roche, Switzerland) and a third-generation Lentiviral Packaging Mix (Merck, Germany), according to the manufacturer’s protocols. In brief, a transfection mixture containing 2 μg of the pSIN-GFP-Fg plasmid (Addgene) was added to HEK293T packaging cells and then incubated at room temperature (RT) for 20 min. Subsequently, 100 μL of the lentivirus-containing supernatant was added dropwise to the organoid-generating cells pre-conditioned with 8 μg/mL polybrene. After 48h, the culture medium was replaced with fresh media containing 5 μg/mL puromycin (Thermo-Fisher, USA). Antibiotic-resistant cells were then selected using a FACSAria IIb cell sorting system (BD Biosciences, USA).

### 3D organoid culture in hydrogels

For the generation of 3D organoids, PDAC or PDAC-GFP organoid-generating cells were grown as described in the “Cell culture” section. In brief, organoid-generating were embedded in non-gelated M, CM, or C mixtures at a final concentration of 3×10^5^ cells/mL. 20 μL of this cell-hydrogel mixture was added to each hydrogel-loading well and left to gelate, as detailed in the “Hydrogel preparation” section. After gelation, regular or TGFβ media was added, and the 3D culture was incubated at 37°C and 5% CO_2_, as required. At this stage, the loaded devices contain EOSs, which consist of small clusters of organoid-generating cells. Experiments requiring mature organoids were conducted after 7 days of incubation.

### Time-lapse microscopy of PDAC “organoid seeds”

Time-lapse movies of GFP-embedded organoid-generating cells were acquired using a Zeiss LSM-880 AxioObserver inverted CLSM equipped with a 10x Plan-Neofluar (0.30 NA) objective and an excitation wavelength of 488 nm. Immediately after the hydrogel gelation, Z-stacks encompassing a volume of 850×850×150 µm^3^ were acquired every 20 min throughout 16h. Throughout the entire experiment, EOSs were incubated at 37°C and 5% CO_2_.

### Organoid seed tracking

3D EOSs migration tracks were obtained from time-lapse movies processed using the TrackMate v6.0.3 single-particle tracking plugin (Tinevez et al., 2017) for Fiji. The TrackMate particle-tracking parameters used for 3D EOSs trajectory reconstruction are summarized in **Supplementary Table 2**. The mean speed (μm/h) (Equation 1) was calculated using a homemade script for Python. Additionally, the directness index was quantified by dividing the Euclidean distance (μm) by the accumulated distance (µm) between the starting and endpoint of a migrating EOS. A directness value close to 0 indicates indirect migration, while a value close to 1 suggests straight-line migration. At least 400 organoid seeds were analyzed across three independent experiments.

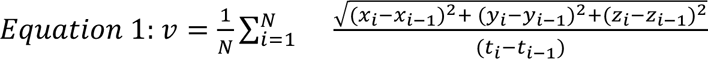

### Quantification of cell protrusions in organoid seeds

Protrusions from EOSs were analyzed using a homemade Fiji script. Time-lapse movies were processed using the CLIJ2 package for Fiji (Haase et al., 2020). First, 3D videos were transformed into 2D time-series images using a MIP projection followed by a Laplacian of Gaussian (LoG) filtering for edge detection. Then, EOSs were segmented using the Weka Segmentation (Arganda-Carreras et al., 2017) plugin for Fiji, generating a binary mask. Next, binary morphological opening steps were performed to remove cell protrusions. Finally, the logical operator XOR was applied to obtain the protrusion mask. Subsequently, the number and length of segmented protrusions were analyzed using MorpholibJ (Legland et al., 2016). A minimum of 250 protrusions were analyzed across three independent experiments.

### Quantification of organoid morphology

PDAC-GFP organoids were grown as described in the “3D organoid cultures in hydrogels” section. On day 7, samples were fixed with 4% paraformaldehyde (PFA) at 37°C for 30 min, washed thoroughly, and cell nuclei were stained with SiRDNA far-red labeling probe (Spirochrome, USA) for 1h at RT.

Image acquisition was performed using a Zeiss LSM-880 AxioObserver inverted CLSM equipped with a 25x LD LCI Plan-Aprochomat (0.8 NA, W) objective. Image stacks with a total sample volume of 1065×1065×200 μm^3^ were acquired using a Mai-Tai® DeepSee™ Ti-Sapphire laser. The laser was sequentially set to 740 nm for scanning nuclei and 920 nm for scanning cytoplasmic GFP.

Prior to quantification, nuclei and cytoplasmic were segmented using StarDist 3D (Schmidt et al., 2018) and the Weka Segmentation 3D (Arganda-Carreras et al., 2017) plugin, respectively. The StarDist 3D model was trained from scratch for 50 epochs on 45 paired image patches (patch size: (72,72,32), batch size: 1, number of rays: 32, augmentation: true), accelerated using an NVIDIA Quadro P1000 GPU on an Intel^(R)^ Core^(TM)^ i7-8700K CPU @ 3.70GHz 64GB RAM Windows 10 Pro-machine. Then, the generated masks were analyzed using a homemade Fiji script. In brief, cytoplasmic masks were pre-processed using GPU-accelerated 3D median filter and binary closing available in the CLIJ2 library (Haase et al., 2020). Then, each organoid was labeled, and the intersection between the cytoplasmic mask and the nuclei mask was calculated using a logic AND operator. A set of morphological descriptors was quantified using MorpholibJ (see **Supplementary Table 3**). A minimum of 150 organoids were analyzed across three independent experiments.

### Classification of PDAC organoid morphology

PDAC organoid morphology classification was conducted using a support vector machine (SVM) learning algorithm implemented via Scikit-Learn (Pedregosa et al., 2012) Python machine learning library. This model differentiated between organoid phenotypes based on the morphological descriptors previously extracted in the “Quantification of Organoid Morphology” section. The SVM structure employed a radial basis function (RBF) kernel and both the regularization parameter (set to *C* = 1) and the γ value internally calculated by Scikit-Learn. Moreover, according to the morphological descriptors, each feature was standardized using z-score normalization. Finally, the most relevant morphological features were selected based on their highest κ-statistic scores (see **Supplementary Table 3**), allowing for dimensionality reduction of the dataset.

Two separate SVM models were trained for each media conditioning (regular and TGFβ), using an independent dataset annotated by an expert from image stacks obtained in the “Quantification of Organoid Morphology” section. The SVM trained for regular-conditioned data classified three morphologies: Cyst, Solid, and Invasive. The SVM model for TGFβ-conditioned data functioned as a binary classifier, distinguishing between Solid and Invasive organoid morphologies. The decision to use two different SVMs was driven by the observed morphological differences between the conditions prior to organoid annotation, as cystic forms were absent in the TGFβ setting. Model validation via 10-fold cross-validation achieved accuracies of 95% for regular media and 100% for TGFβ media.

Additionally, the distribution of organoid morphologies was visualized using a t-SNE plot, implemented with Scikit-Learn. The t-SNE models were configured for 1,000 iterations with a perplexity of 50, employing an Euclidean metric to calculate distances between morphological descriptors after organoid classification. Specifically, two t-SNE models were created from the result of each SVM classifier to visualize the distribution of morphologies of organoids grown in regular and TGFβ-conditioned media, respectively. A third t-SNE model was produced by combining the results of both SVM classifiers to visualize the distribution of morphologies, regardless of the cultured media used.

### Collagen-I remodeling assays

PDAC organoids were grown as described in the “3D organoid cultures in hydrogels” section. On day 7, organoids were fixed with 4% PFA solution at 37°C for 30 min. Then, cells were counterstained with SiR-DNA and rhodamine-phalloidin (Abcam, USA) for F-actin filament staining, following the manufacturer’s protocols. PDAC-induced collagen-I remodeling was imaged as described in the “Hydrogel Morphological Characterization” section.

Fiber anisotropy (α) and compaction were analyzed following the method described previously by our group (Anguiano et al., 2020). Twelve images were analyzed across three independent experiments.

### qRT-PCR assays

Total RNA from mature organoids was extracted using a Maxwell RSC SimplyRNA Kit (Promega, Spain) following the manufacturer’s protocol. Reverse transcription was performed using M-MLV reverse transcriptase, and quantitative RT-PCR (qRT-PCR) was conducted with SYBR Green PCR Master Mix (Thermo-Fisher, USA) on a QuantStudio3 system (Thermo-Fisher, USA). The PCR master mix included the primers described in **Supplementary Table 4**.

Relative expression of each transcript was calculated according to this formula 2^−ΔΔCT^ = [(CT _gene of interest_ - CT _housekeeping_) Organoids - (CT _gene of interest_ - CT _housekeeping_) 2D-platted cells], where CT corresponds to cycle number (Schmittgen & Livak, 2008). Expression data were normalized to endogenous GAPDH levels. The fingerprint signature was visualized as a hierarchical heat-map using a simple Python script (v.3.12.6) that uses the Seaborn library (Waskom, 2021). Six independent experiments were performed per hydrogel type.

### Immunofluorescence labeling

PDAC organoids were grown as described in the “3D Organoid Cultures in Hydrogels” section. On day 7, organoids were fixed with 4% PFA at 37°C for 30 min and then permeabilized with 0.1% Triton-X at RT for 15 min. Non-specific interactions were blocked by incubating the samples in a 1% BSA at RT for 30 min. Subsequently, the samples were labeled with the specific EMT antibodies listed in **Supplementary Table 5**, followed by incubation at 4°C for 48h.

Specifically, live mitochondria labeling was performed by incubating the organoids with regular media containing 5 μmol/L of the MitoTracker™ Green probe (Thermo-Fisher, USA) for 20 min at 37°C.

Z-stack CLSM images were acquired using a Zeiss LSM-800 AxioObserver inverted CLSM equipped with a 40x Plan-Aprochromat (1.2 NA, W), adjusting the excitation wavelength as required by the spectral properties of the conjugated antibodies.

### *CDH1* and CTNNB1 distribution analysis

The distribution of *CDH1* and *CTNNB1* was analyzed using a Fiji script adapted from Nyga et al. (Nyga et al., 2021) for Z-stack imaging. In brief, an ROI encompassing at least two nuclei was manually selected. For CDH1 or CTNNB1 quantification at the cell junction, a line was drawn connecting two nuclei. The cytoplasmic CDH1 intensity was quantified by calculating the average pixel intensity above Otsu’s threshold within the sample volume(Otsu, 1979). Then, the junctional and sample volume CDH1 masks were intersected using the logical AND operator. In parallel, the SiRDNA signal was segmented using Otsu’s threshold, and the average CTNNB1 intensity within the intersecting nuclei mask was determined using the logical AND operator.

The junctional to cytoplasmic CDH1 intensity ratio was calculated by dividing the junctional intensity by the average cytoplasmic intensity. Similarly, the junctional to nuclear CTNNB1 intensity ratio was obtained by dividing the junctional intensity by the average nuclear intensity. Fifty cell-to-cell junctions were analyzed across three independent experiments.

### *CDH1*/*VIM* ratio analysis

The cytoplasmic intensity of the *VIM* channel was determined by calculating the average pixel intensity using Otsu’s threshold method, as described above. The *CDH1*/*VIM* ratio was then calculated by dividing the average junctional *CDH1* value by the average cytoplasmic *VIM* value, followed by a log transformation. This ratio classifies PDAC organoids into epithelial (log *CDH1*/*VIM* ratio > 0) or mesenchymal (log *CDH1*/VIM ratio < 0) phenotypes (El Amrani et al., 2019b). Fifty cell-to-cell junctions were analyzed across three independent experiments.

### Mitochondria morphology analysis

Mitochondrial mass (weighted per cell number), mean volume, sphericity, and branches were quantified using the “3D Mitochondria Analyzer” plugin developed for Fiji (Schindelin et al., 2012b) with default parameters. A minimum of 300 cells per hydrogel type were analyzed from three independent experiments.

### Seahorse assays

Organoid metabolism was analyzed using an XF24 Extracellular Flux Analyzer (Agilent Technologies, Spain). In brief, PDAC organoids were cultured as described in the “3D Organoid Cultures in Hydrogels” section. On day 7, fresh media containing non-buffered DMEM with 50 mmol/L glucose, 4 mmol/L Na-Pyruvate, and 2 mmol/L glutamine (all from Merck, Germany) was added. Oxygen consumption rates (OCR) and extracellular acidification rates (ECAR) were measured under baseline conditions and following the addition of 0.5 μmol/L oligomycin, 1 μmol/L FCCP, 1 μmol/L antimycin-A, 1 μmol/L rotenone (all from Merck, Germany), and 100 mM 2-deoxy-d-glucose (2-DG).

Spare respiratory capacity (SRC) and glycolytic capacity (GC) were calculated as the differences between their maximum and basal values. Mitochondrial- and glycolytic-ATP production rates were calculated as described by Desousa et al. (Desousa et al., 2023). Data was normalized to the µg of protein of each well. Five independent experiments were performed per condition.

### Flow cytometry

Organoid mitochondrial membrane potential (Δψm) was analyzed by flow cytometry using a MitoProbe-TMRM Assay Kit (Thermo-Fisher, USA). On day 7, organoids were isolated from the matrices and disaggregated into individual cells. Live mitochondria were stained by incubating cells in fresh media containing 125 ng/ml TMRM dye for 20 min at 37°C. Subsequently, samples were thoroughly washed and collected using a CytoFLEX system (Beckman Coulter, USA). Data analysis was performed using FlowJo software (TreeStar, USA). Three independent experiments were analyzed per condition.

### *In vitro* live/dead assays

Organoid sensitivity to Gem (MedChemExpress, Sweden) was assessed using a live/dead assay kit (Thermo-Fisher, USA). In brief, after 5 days of incubation, the medium was replaced with 100 µL of Gem solution at a final concentration of 0,1 µM, followed by a treatment period of 48h at 37°C and 5% CO_2_. On day 7, samples were thoroughly rinsed, and Syto9 (green, live) and propidium iodide (PI) (red, dead) were added to a final concentration of 7 µM and 40 µM, respectively, followed by a 2h incubation period. The IG_50_ value for Gem was determined using PDAC organoids embedded in Matrigel and cultured in regular media with the following Gem concentrations: 0, 0.001, 0.1, 1, 10, 100, and 1000 µM.

Organoids were imaged using a Zeiss Cell Observer Spinning Disk microscope equipped with a 25x LD LCI Plan-Apochromat (0.8 NA, W) objective. At least 10 images comprising a total sample volume of approximately 607×607×150 µm^3^ were acquired. Image acquisition was performed using an excitation wavelength of 488 nm for Syto9 (live) visualization and 561 nm for PI (dead) visualization, respectively.

Viability quantification was measured using a custom script in Fiji. In brief, Syto9 and PI signals were segmented separately using fixed-intensity threshold values. The PI and Syto9 masks were intersected using the logical AND operator to generate the live organoid mask. Relative viability was calculated as the ratio between Gem-treated and regular media conditions based on the live organoid masks. Ten images containing at least 100 organoids were analyzed across three independent experiments.

### Mice handling and subcutaneous model

Male C57BL/6J wild-type mice (Charles River, France), aged 5-6 weeks, were housed under pathogen-free conditions at the CIMA animal facility (registration number ES31-2010000132). Animal handling and procedures were conducted under a protocol approved by the Animal Ethics Committee of the University of Navarra (Study 029-22).

To establish a syngeneic PDAC model, organoids cultured as described in the “3D Organoid Cultures in Hydrogels” section were subcutaneously injected into the lower right flank on day 7, preserving the embedding matrix (M, CM, and C). When the tumor volume reached 50 mm^3^, as measured by echography on day 14, the mice were randomly assigned to six experimental groups (n = 9 per group). They were treated with either vehicle control (0.9% saline, i.p.) or metronomic gemcitabine (100 mg/kg, i.p., twice a week for two weeks). On day 28, tumor volume and vasculature were scanned using the Vevo3100 ultrasound system (VisualSonics, Canada) coupled to an MX550D transducer. Mice were anesthetized with a 2% isoflurane/air mixture (Isoflo^®^, ABBOTT, Spain) and positioned on a heated pad. ECG sensors were attached to the limbs for comprehensive vital sign monitoring. Once tumors were located, B-mode images, combined with color Doppler-mode, were automatically acquired along the entire tumor length. Tumor volume (weighted per the μg of organoids inoculated) and vasculature (weighted per tumor volume) were quantified using VevoLab software (v5.8.1, VisualSonics, Canada). For this purpose, a trained technician semi-automatically delineated the tumor margins. Relative tumor growth inhibition (TGI) was calculated using the formula: TGI = (Vf - Vi)/Vi, where Vi and Vf are the tumor sizes on days 14 and 28, respectively. A TGI > 0 indicates tumor growth, while a TGI < 0 indicates tumor regression. Subcutaneous tumors were harvested on day 28.

### Statistical Analysis

Each experiment was repeated at least three times. The normality of data distribution was assessed using Kolmogorov-Smirnov and Shapiro-Wilk tests. Data from all conditions were required to pass the normality test to be included in parametric testing: (i) for parametric data, a T-test analysis was performed between two independent samples. For comparisons involving more than two groups, analysis of the variance (ANOVA) was performed, followed by Tukeýs or Tamhane’s T2 multiple comparison test, depending on the variance status; (ii) for non-parametric data with more than two groups, the Kruskal-Wallis test was used, followed by Dunn’s multiple comparison test. For comparisons between two independent samples, the Mann-Whitney U test was applied. All statistical analyses were performed using GraphPad Prism-8 (GraphPad, USA). Datasets were presented as scatter plots, displaying the mean ± standard deviation.

