## Supplementary Information file for "Quantitative evaluation of the role of the microenvironment in the phenotype, metabolism, and drug resistance of PDAC tumor organoids"

### Supplementary Tables:

**Supplementary Table 1**. Principal reagents used to supplement the organoid culture media.

| REAGENT | SOURCE | CONCENTRATION | OBSERVATIONS |
|---|---|---|---|
| **Advanced DMEM F12** | Gibco | 1X, (base medium) | Regular media/TGFβ media |
| **Penicillin/Streptomycin** | Gibco | 1X | Regular media/TGFβ media |
| **HEPES** | Lonza | 1X | Regular media/TGFβ media |
| **GlutaMax** | Gibco | 1X | Regular media/TGFβ media |
| **A83-01** | TOCRIS | 0.5μM | Regular media |
| **mEGF** | Life Technologies | 50 ng/mL | Regular media/TGFβ media |
| **hFGF10** | Peprotech | 100 ng/mL | Regular media/TGFβ media |
| **Gastrin I** | TOCRIS | 10 nM | Regular media/TGFβ media |
| **mNoggin** | Peprotech | 100 ng/mL | Regular media |
| **Y-27632** | Sigma | 14 μM | Regular media/TGFβ media |
| **N-acetylcysteine** | Sigma | 1.25 mM | Regular media/TGFβ media |
| **Nicotinamide** | Sigma | 10 mM | Regular media/TGFβ media |
| **B-27** | Life Technologies | 1X | Regular media/TGFβ media |
| **TGFβ** | R&D Systems | 1 ng/mL | TGFβ media |

**Supplementary Table 2**. TrackMate particle-tracking parameters used for 3D organoid trajectory reconstruction.

| PROCESSING STEP | PARAMETER/VALUE |
|---|---|
| LoG detector | radius = 12.5 μm<br>threshold = 2.5 μm<br>subpixel = True<br>median = True |
| Simple LAP tracker | max_frame_gap = 0<br>max_distance = 20.0 μm<br>max_gap_distance = 20.0 μm |

**Supplementary Table 3**. Morphological descriptors used for PDAC organoids characterization using MorpholibJ and their respective κ-statistic scores.

| FEATURE | DESCRIPTION | F-Score |
|---|---|---|
| **Circularity:** | The normalized ratio of the area over the square of the perimeter ($4\pi \cdot A/p^2$) | 180.83 |
| **Convexity:** | The ratio of the perimeter of the convex hull of the particle over the perimeter of the particle | 173.87 |
| **Sphericity:** | The ratio of the squared volume over the cube of the surface area ($36\pi V^2/S^3$). Normalized so that the value for a perfect sphere equals one. | 87.72 |
| **Geodesic.Elong:** | The ratio of geodesic diameter over the diameter of the largest inscribed circle | 87.47 |
| **R.Lumen:** | The radius of the largest inscribed sphere for the lumen of an organoid | 58.12 |
| **R1.Ellipse:** | The radius of the major axis of an ellipse | 45.59 |
| **Ellipse.Elong:** | The ratio of the largest over the smallest axis lengths | 41.00 |
| **Vol Lumen/Vol Total:** | The ratio between the total number of voxels comprising the *Vol Lumen* and *Vol Total* | 39.92 |
| **GeodesicDiameter:** | The length of the longest geodesic path within a particle | 38.64 |
| **R1:** | The length of the radius 1 of an ellipsoid | 34.51 |
| **Ball Radius:** | The minimum sphere circumscribed in the *Vol Total* | 29.95 |
| **InscrDisc.Radius:** | The minimum radius inscribed in the *Vol Total* | 28.07 |
| **R.Organoid:** | The radius of the largest inscribed sphere within an organoid | 27.88 |
| **R1/R3:** | The elongation of the object defined as the ratio between the length and width | 27.65 |
| **R2:** | The length of the radius 2 of an ellipsoid | 26.48 |
| **R3:** | The length of the radius 3 of a 3D ellipsoid | 26.28 |
| **R2.Ellipse:** | The radius of the minor axis of an ellipse | 25.35 |
| **Perimeter:** | Length of the boundary comprising a particle, using the Crofton formula. | 24.19 |
| **Nuclei Number:** | The number of nuclei corresponding to the instances obtained from SirDNA Staining | 15.89 |
| **Surface GFP:** | The total area of the uppermost layer of the organoid GFP instance. | 12.91 |
| **ConvexArea:** | The minimum enclosed convex-shape area (convex hull) of a particle | 11.59 |

| | | |
|---|---|---|
| **Organoid Area:** | The number of pixels comprising the organoid instance multiplied by the area of each pixel applied to the maximal projection of the organoid mask. | 11.39 |
| **Vol GFP:** | The number of voxels comprising the GFP staining | 10.64 |
| **Vol Total:** | The total number of voxels obtained by the sum of *Vol GF*P and *Vol Lumen* | 9.84 |
| **R2/R3:** | The flatness of the object defined as the ratio between the width and height | 7.08 |
| **Vol Lumen:** | The number of voxels comprising the organoid lumen | 4.69 |
| **Tortuosity:** | The ratio between the shortest pathway to the distance between the inlet and outlet plane | 4.56 |

**Supplementary Table 4.** List of EMT PCR primers.

| **GENE** | **MARKER-FAMILY** | **SEQUENCE** |
|---|---|---|
| **CDH1-F** | EMT | 5′-AGCGGCAAGAGTGAGATTTCT-3′ |
| **CDH1-R** | EMT | 5′-CCTCCAGGTTATTCTCCAGGG-3′ |
| **CDH2-F** | EMT | 5′-TGAAACGGCGGGATAAAGAG-3′ |
| **CDH2-R** | EMT | 5′-GGCTCCACAGTATCTGGTTG-3′ |
| **VIM-F** | EMT | 5′-CAAGAGCGCCTTGACGATACA-3′ |
| **VIM-R** | EMT | 5′-CCAAGAGACAGGTTTCTCCATC-3′ |
| **ZEB1-F** | EMT | 5′-CTCTGCAAGAGACTTCCATCCAGT-3′ |
| **ZEB1-R** | EMT | 5′-GAAGTAGGGAAGGCCGTGG-3′ |
| **GAPDH-F** | Endogenous | 5′-ACTTTGTCAAGCTCATTTCC-3′ |
| **GAPDH-R** | Endogenous | 5′-TGCAGCGAACTTTATTGATG-3′ |

**Supplementary Table 5**. Conjugated antibodies and concentrations used for the immunofluorescence assays.

| **REAGENT** | **SOURCE** | **CONCENTRATION** |
|---|---|---|
| DAPI | ThermoFisher | 1:400 |
| Anti-E-Cadherin (*CDH1*) (Alexa488) | BD Biosciences | 1:200 |
| Anti β-Catenin (*CTNNB1*) (PE) | R&D Systems | 1:200 |
| Anti-VIM (Alexa647) | Thermofisher | 1:200 |

### Supplementary Figures:

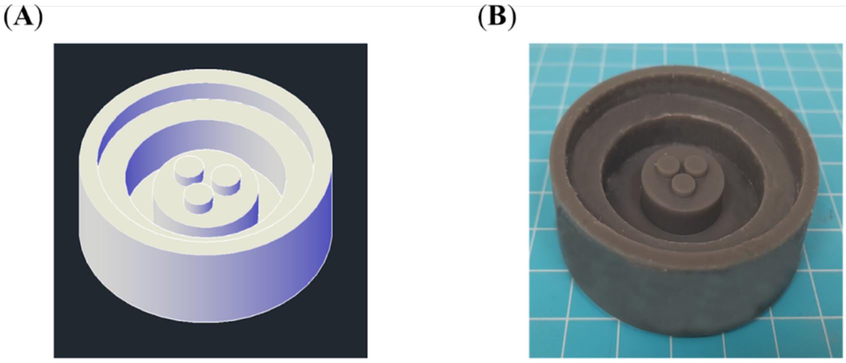


**Supplementary Figure 1.** (**A**), 3D CAD model of the master mold. (**B**), Image of a 3D-printed resin mold created through stereolithography.

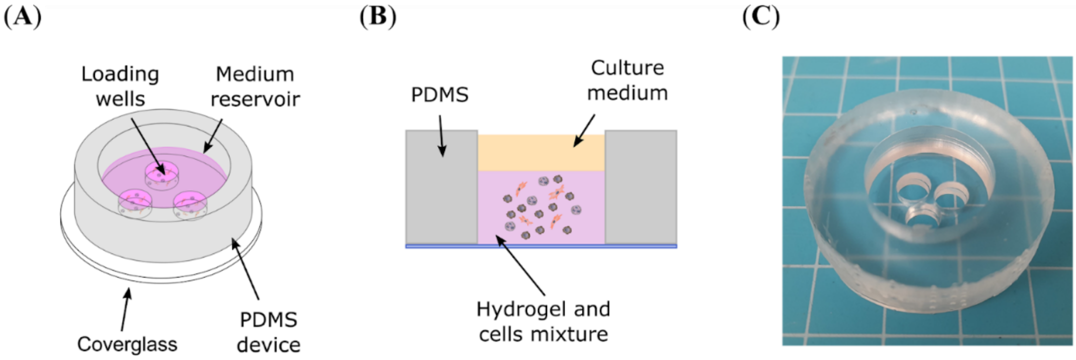


**Supplementary Figure 2.** (**A**), Schematic representation of the micro-device. (**B**), A cross-sectional view illustrating a single well, with organoids embedded within the hydrogel and covered on top with culture media. (**C**), Image of the PDMS micro-device used for PDAC organoid culture.

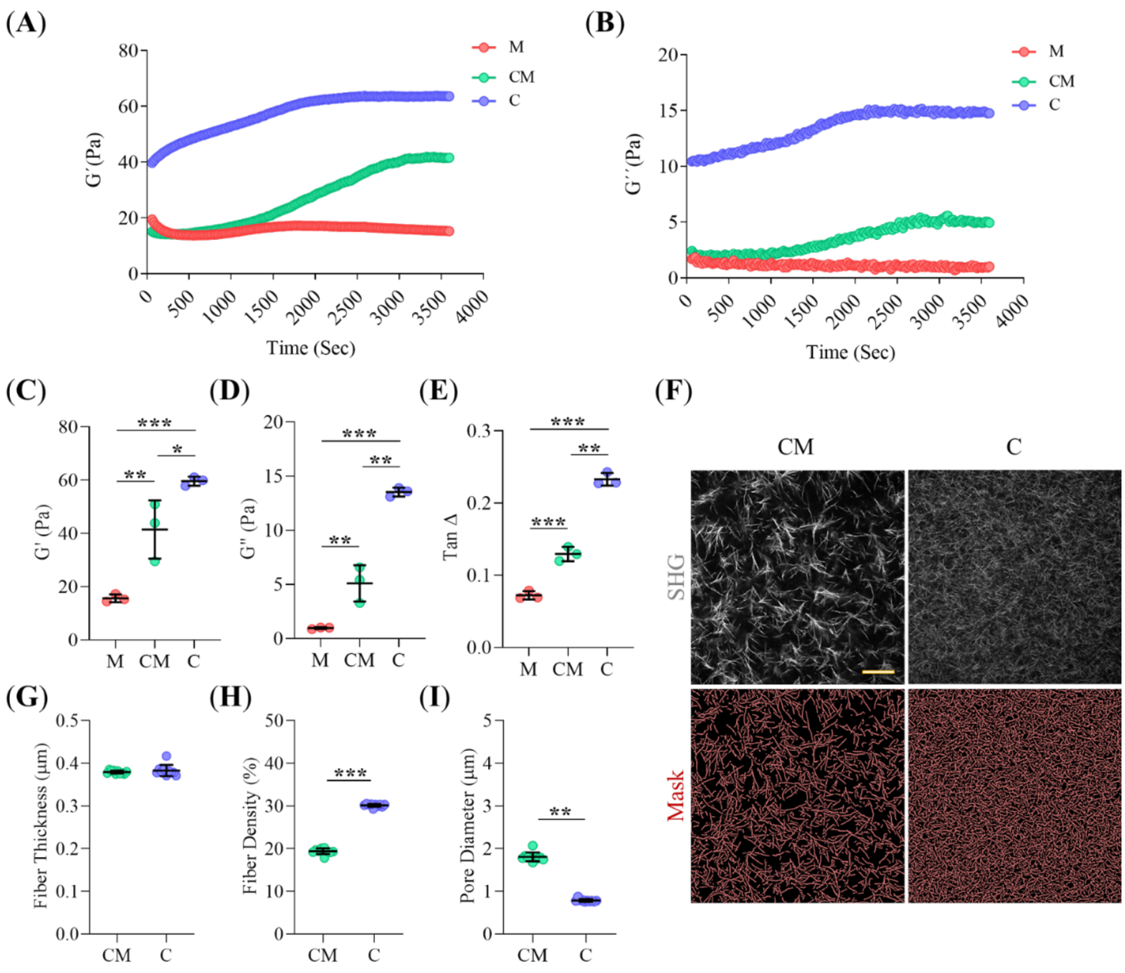


**Supplementary Figure 3.** (**A-B**), Storage (G', Pa) and loss (G", Pa) moduli profiles during oscillatory time sweep assays for M, CM, and C hydrogels. (**C-E**), Quantification of storage (G') and loss (G") moduli, and G'/ G" ratio (TanΔ) of M, CM, and C hydrogels after gel polymerization, extracted from oscillatory time sweep assays. (**F**), Z-stack projections of representative 3D second harmonic MP-CLSM images of Collagen-I fiber lattice in CM and C hydrogels. Segmentation masks (in red) depict the architecture of the Collagen-I lattice. Scale bars represent 15 µm. (**G-I**), Quantitative analysis of Collagen-I fiber lattice morphology, including fiber thickness (µm), fiber density (%), and pore diameter (µm), respectively, based on the images shown in F. ** Indicates highly statistically significant differences between groups (p<0.001). ** Indicates moderately statistically significant differences between groups (p<0.01). * Indicates statistically significant differences (p<0.05).

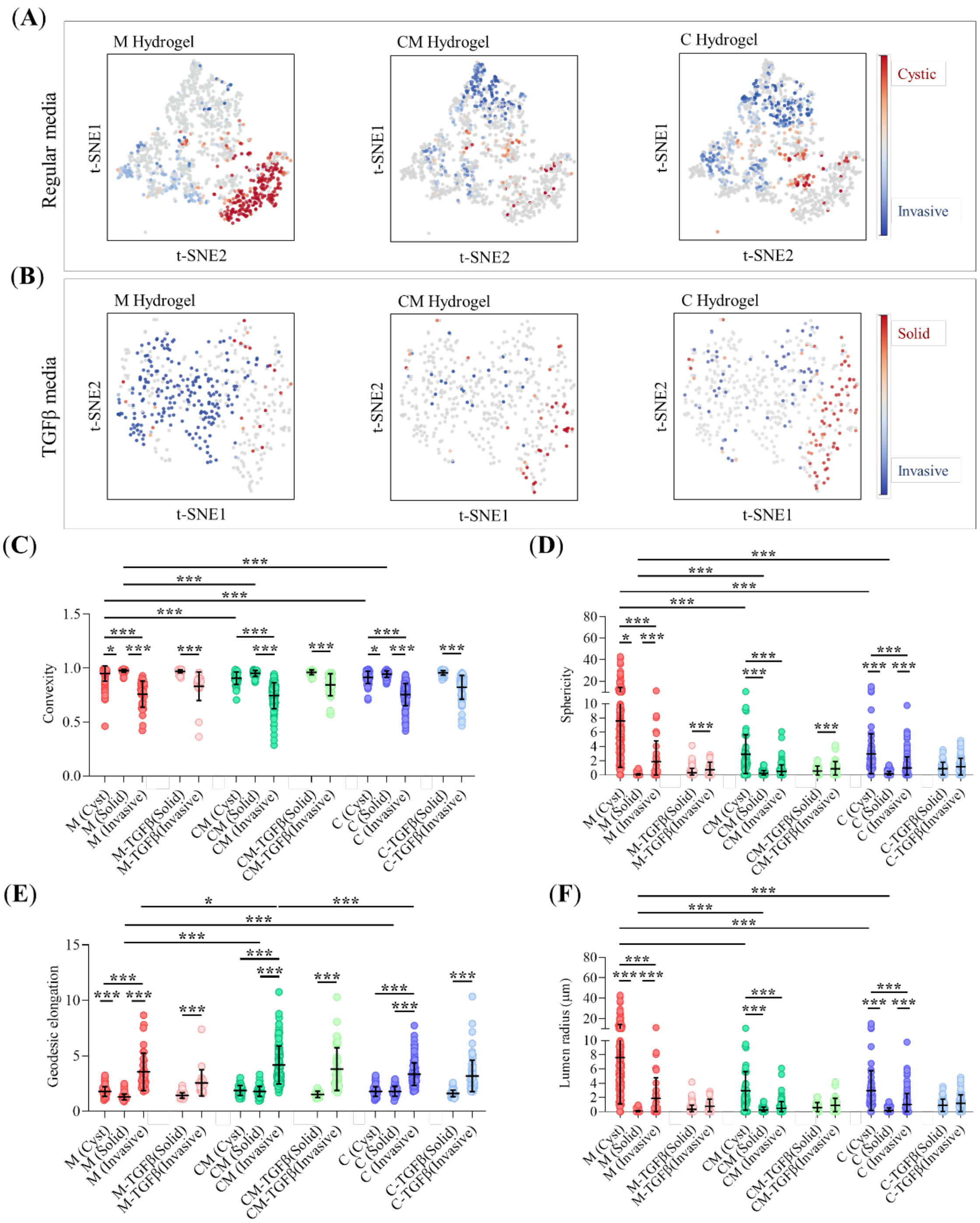


**Supplementary Figure 4.** (**A-B**), t-SNE projections illustrating the distribution of morphological classes for PDAC organoids under regular and TGFβ conditioning, respectively, categorized by gel type (M, CM, and C hydrogels). The scale bar represents the probability of classification as Cyst, Solid, or Invasive. (**C-F**), Quantification of the highest κ-scored morphological descriptors used in the SVM classifiers, including organoid convexity, geodesic elongation, and lumen radius, categorized by morphological classes for each gel type and cell media. $^{***}$ Indicates highly statistically significant differences between groups ($p<0.001$). $^{*}$ Indicates statistically significant differences between groups ($p<0.05$).

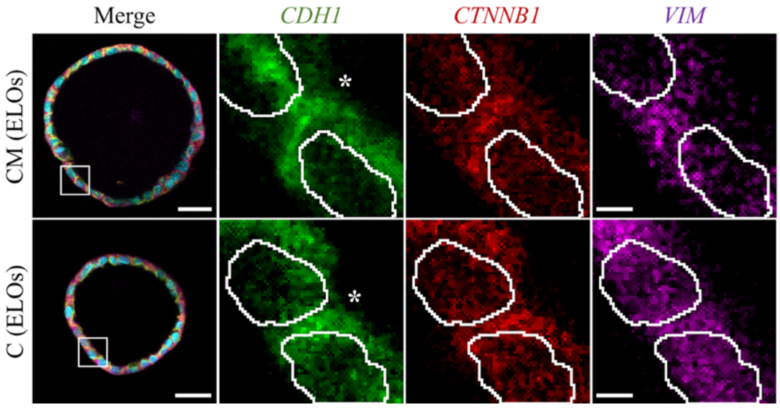


**Supplementary figure 5.** (**A**) Z-stack projections of representative CLSM images in the different PDAC organoid classes cultured in CM and C hydrogels under regular or TFGβ conditioning, respectively. Staining: E-cadherin (*CDH1*, green), β-catenin (*CTNNB1*, red), vimentin (*VIM*, magenta), and nuclei (SiRDNA, cyan). Scale bars 30 µm. White square insets provide a zoomed-in view of cell-cell junctions. White asterisks indicate the location of CDH1 at the cell-cell junctions. Scale bars: 3 µm.
